# Intense light-mediated circadian cardioprotection via transcriptional reprogramming of the endothelium

**DOI:** 10.1101/561340

**Authors:** Yoshimasa Oyama, Colleen M. Bartman, Stephanie Bonney, J. Scott Lee, Lori A. Walker, Jun Han, Christoph H. Borchers, Peter M. Buttrick, Carol M. Aherne, Nathan Clendenen, Sean P. Colgan, Tobias Eckle

**Affiliations:** Mucosal Inflammation Program, Departments of Medicine and Anesthesiology, University of Colorado Anschutz Medical Campus, Aurora, CO, USA; Division of Cardiology, Department of Medicine, University of Colorado Anschutz Medical Campus, Aurora, CO, USA; Department of Biochemistry and Microbiology, Genome BC Proteomics Centre, University of Victoria, Victoria, BC, Canada; Graduate Training Program in Cell Biology, Stem Cells, and Development, University of Colorado Anschutz Medical Campus, Aurora, CO, USA

**Author notes:** authors contributed equally (Yoshimasa Oyama, Colleen M. Bartman, Stephanie Bonney). Lead Contact*: Tobias Eckle, M.D., Ph.D. Correspondence should be addressed to, Tobias Eckle, M.D., Ph.D., Professor of Anesthesiology, Cardiology and Cell Biology Department of Anesthesiology, University of Colorado Denver, 12700 E 19th Avenue, Mailstop B112, RC 2, Room 7121, Aurora, CO 80045; or - 2947.

## Abstract

Consistent daylight oscillations and abundant oxygen availability are fundamental to human health. Here, we investigate the intersection between light-(Period 2, PER2) and oxygen-(hypoxia inducible factor, HIF1A) sensing pathways in cellular adaptation to myocardial ischemia. We demonstrate that intense light is cardioprotective via circadian PER2 amplitude enhancement, mimicking hypoxia elicited adenosine- and HIF1A-metabolic adaptation to myocardial ischemia under normoxic conditions. Whole-genome array from intense light exposed wildtype or *Per2^-/-^* mice and myocardial ischemia in endothelial-specific PER2 deficient mice uncover a critical role for intense light in maintaining endothelial barrier function via light-enhanced HIF1A transcription. A proteomics screen in human endothelia reveals a dominant role for PER2 in metabolic reprogramming to hypoxia via mitochondrial translocation, TCA cycle enzyme activity regulation and HIF1A transcriptional adaption to hypoxia. Translational investigation of intense light in human subjects identifies similar PER2 mechanisms, implicating the use of intense light for the treatment of cardiovascular disease.

**GRAPHICAL ABSTRACT:** 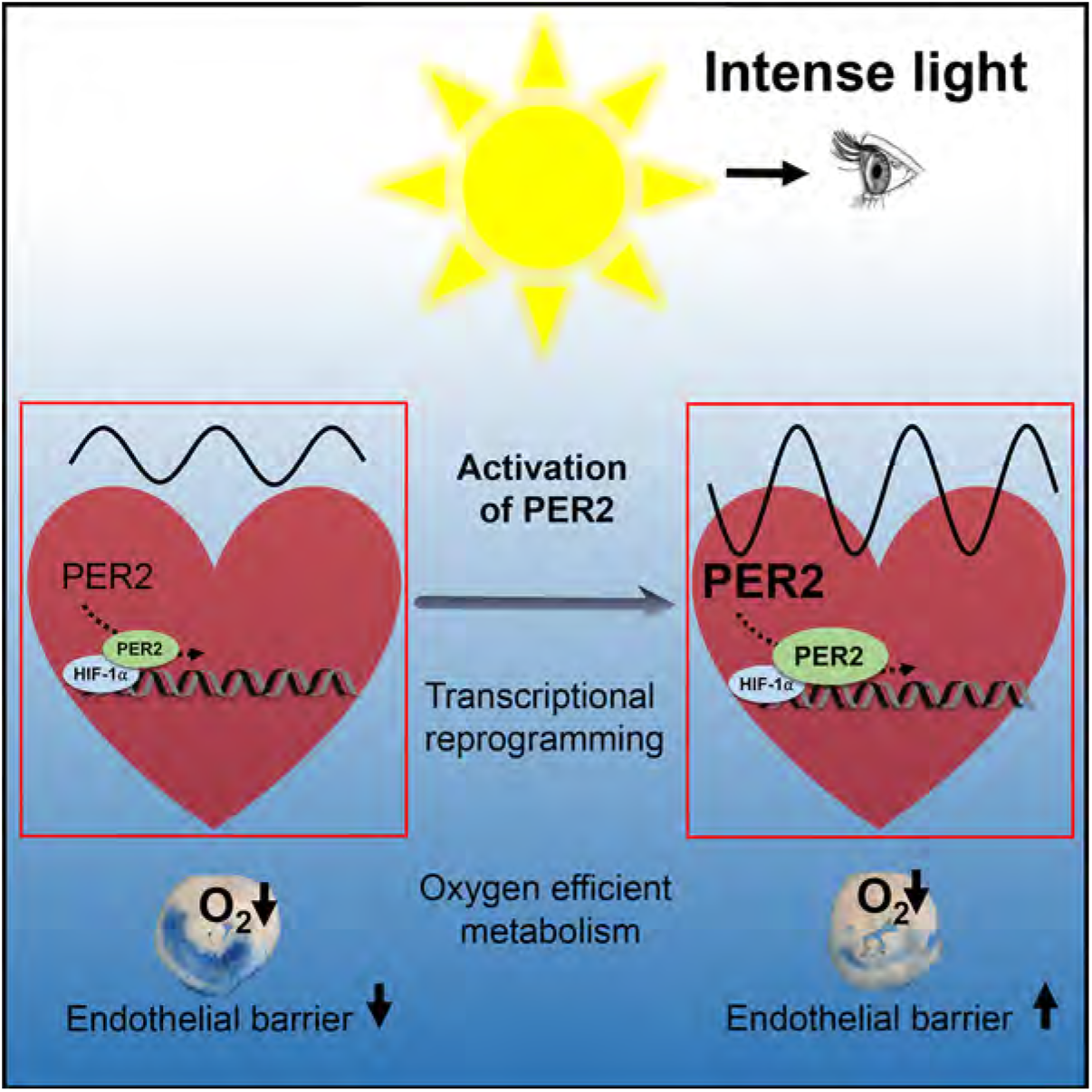

## INTRODUCTION

The appearance of sunlight and oxygen on earth were undoubtedly the most dramatic environmental changes during evolution (Zerkle et al., 2017). As a result, almost all organisms on this planet are equipped with light- and oxygen-sensing pathways. Notably, light sensing and oxygen sensing pathways are linked on a cellular level in mammals (Gu et al., 2000; Hogenesch et al., 1998; McIntosh et al., 2010). Hypoxia inducible factor 1α (HIF1A), an evolutionarily conserved transcription factor enabling cellular adaptation to low oxygen availability (Semenza, 2011), belongs to the same protein family as the light-inducible circadian core protein Period 2 (PER2) (Liu et al., 2012). Both belong to the *PAS* domain superfamily of signal sensors for oxygen, light, or metabolism (Hogenesch et al., 1998; Taylor and Zhulin, 1999). As such, *Hif1*α mRNA levels cycle in a circadian manner in mouse cardiac tissue (Eckle et al., 2012) and rhythmic oxygen levels reset the circadian clock through HIF1A (Adamovich et al., 2017). This evolutionarily conserved relationship between light (circadian)- and oxygen-sensing pathways suggest a role for light elicited circadian rhythm proteins in disease states of low oxygen availability, such as myocardial ischemia.

In the present studies, we sought to develop a cardioprotective strategy using light to target and manipulate PER2 function and uncover mechanisms of PER2 dependent adaptation to hypoxia or ischemia (Eckle et al., 2012). In a comprehensive and systems biology approach, we dissect light and hypoxia elicited pathways in mice and humans from a cellular level to the whole body. Our investigations reveal that circadian PER2 functions at the crossroads between light elicited circadian amplitude enhancement and transcriptional HIF1A-dependent adaptation to oxygen depletion in hypoxia or ischemia. Combined, we demonstrate a mechanistic understanding of cardioprotection with light therapy by targeting and manipulating hypoxic pathways to reduce infarct sizes after myocardial ischemia.

## RESULTS

### Intense light elicited cardiac PER2 amplitude enhancement as a cardioprotective mechanism

Intense light is the dominant regulator of human circadian rhythms and PER2 activity (Albrecht et al., 2001; Remi, 2015). Here we investigated intense light exposure protocols and found that housing mice under “intense light conditions” (10,000 LUX, full spectrum, UV-filter, L(light):D(dark) phase 14:10h) robustly enhances cardioprotection, reflected as a time-dependent decrease in infarct size and circulating troponin-I levels (Figure 1A-C). Evaluation of locomotor activity during intense light conditions, as determined by wheel running, excluded a phase shift of the circadian period, but identified increases of the total distance walked or the circadian amplitude (Figure 1D-F, Figure S1). Indeed, housing PER2 reporter mice under intense light for one week, revealed increases in the circadian peak and trough levels of cardiac PER2 protein levels (Figure 1G). Further analysis of wheel running activity in *Per2^-/-^* mice demonstrated intense light elicited increases of the total distance walked or the circadian amplitude to be PER2 dependent (Figure 1H-J, Figure S1).

**Figure 1.**
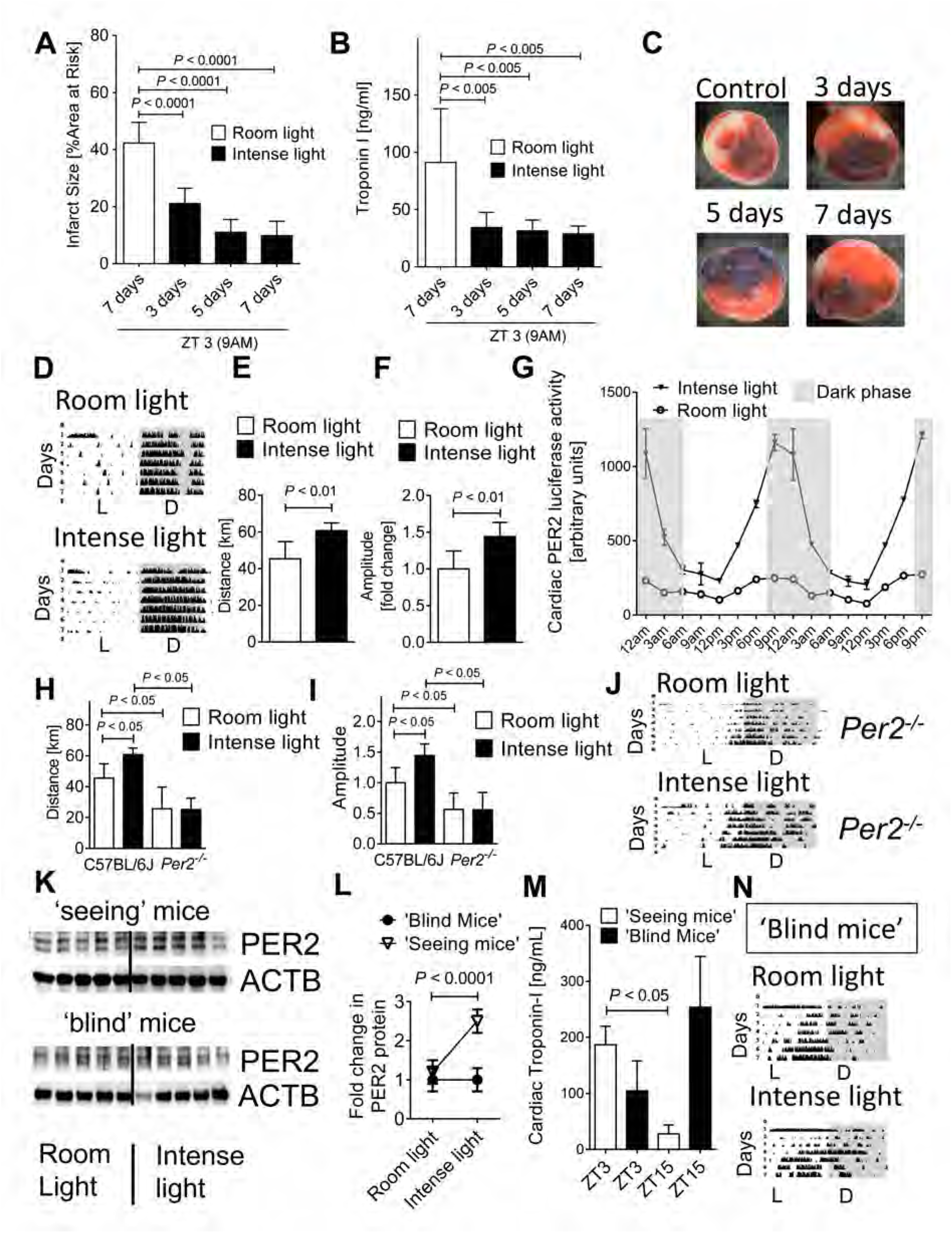
Intense light elicited circadian PER2 amplitude enhancement in cardioprotection. **(A-C)** C57BL/6 mice housed under intense light conditions (IL, 10,000 LUX, L:D 14:10 h) for 3, 5, or 7 days were subjected to 60 min of *in situ* myocardial ischemia followed by 2h reperfusion at ZT3 (9AM) and compared to mice housed under standard room light (RL, 200 LUX, L:D 14:10 h, 7 days); mean±SD; n=6; ANOVA with Tukey’s multiple comparisons test). **(B)** Parallel measurements of serum troponin-I by ELISA (mean±SD; n=6; ANOVA with Tukey’s multiple comparisons test). **(C)** Representative images of infarcts. **(D-F)** Wheel running measurements during 7 days of RL or IL housing in C57BL/6J mice (L=light phase, D=dark phase, n=6; Student’s t test). **(G)** Cardiac PER2 luciferase activity indicating protein in mice after RL or IL for 7 days (mean±SD; n=4, all IL vs RL *P* <0.05, ANOVA with Tukey’s multiple comparisons test). **(H-J)** Wheel running during 7 days of RL or IL housing in C57BL/6J and *Per2^-/-^* mice (n=5-6; ANOVA with Tukey’s multiple comparisons test). (**K,L**) Immunoblot and quantification for PER2 protein in seeing or enucleated (blind) C57BL/6J mice after 7 days of RL or IL at ZT3 (mean±SD; n=5; Student’s t test). **(M**) Troponin-I serum levels in seeing or blind C57BL/6J mice housed under RL conditions followed by 60 min ischemia and 2 h reperfusion at ZT3 or ZT15 (mean±SD; n=4; ANOVA with Tukey’s multiple comparisons test). **(N)** Wheel running measurements during 7 days of RL or IL housing in ‘blind’ C57BL/6J mice (mean±SD; n=4, Student’s t test, see also Figure S1).

To confirm that cardiac circadian PER2 amplitude enhancement requires visual light perception, we enucleated wildtype mice to remove any light sensing structures. Ocular enucleation induced a complete loss of PER2 stabilization in “blind” mice exposed to intense light conditions compared to “seeing” animals. (Figure 1K, L). Myocardial ischemia and reperfusion studies in blind mice under room light housing conditions found ‘shifted’ cardiac troponin kinetics (Troponin ‘blind’: ZT3 [9AM] < ZT15 [9PM] vs Troponin ‘seeing’: ZT3>ZT15) and slightly overall higher troponin levels in ‘blind; mice (‘blind’ vs ‘seeing’ troponin: 168 vs 118 ng/ml, not significant), indicating a lack of circadian synchronization by light (Figure 1M, Figure S1). Indeed, wheel running activity in ‘blind mice’ demonstrated abolished increase of the circadian amplitude similar to *Per2^-/-^* mice (Figure 1N, Figure S1).

To evaluate if intense light-mediated increases of circulating cortisol levels (Oster et al., 2017) or the temperature (Schibler et al., 2015) could have caused the observed circadian amplitude enhancement, we next measured rectal temperatures or plasma cortisol levels (Gong et al., 2015) following 7 days of intense or room light housing. However, we did not observe increases in plasma cortisol levels, or the body temperature in intense light-exposed mice when compared to controls (Figure S1).

Together, these data demonstrate that intense light elicited circadian amplitude is a cardioprotective strategy which requires PER2 and visual light perception.

### Intense light ‘adenosine-preconditions’ and increases HIF1A-HRE binding in the heart before an ischemic insult

Next, we further deciphered the mechanism of intense light elicited cardiac circadian amplitude enhancement and cardioprotection. First, we evaluated the effect of intense light on infarct sizes or circulating troponin-I levels at ZT15, as one week of intense light housing had increased cardiac PER2 protein levels significantly more at ZT15 (9PM) compared to ZT3 (9AM) (Figure 1G). However, we only found slightly smaller infarct sizes or troponin-I levels at ZT15 compared to ZT3 (Figure 2A). Thus, all following studies focused on the robust cardioprotective effect observed at ZT3.

**Figure 2.**
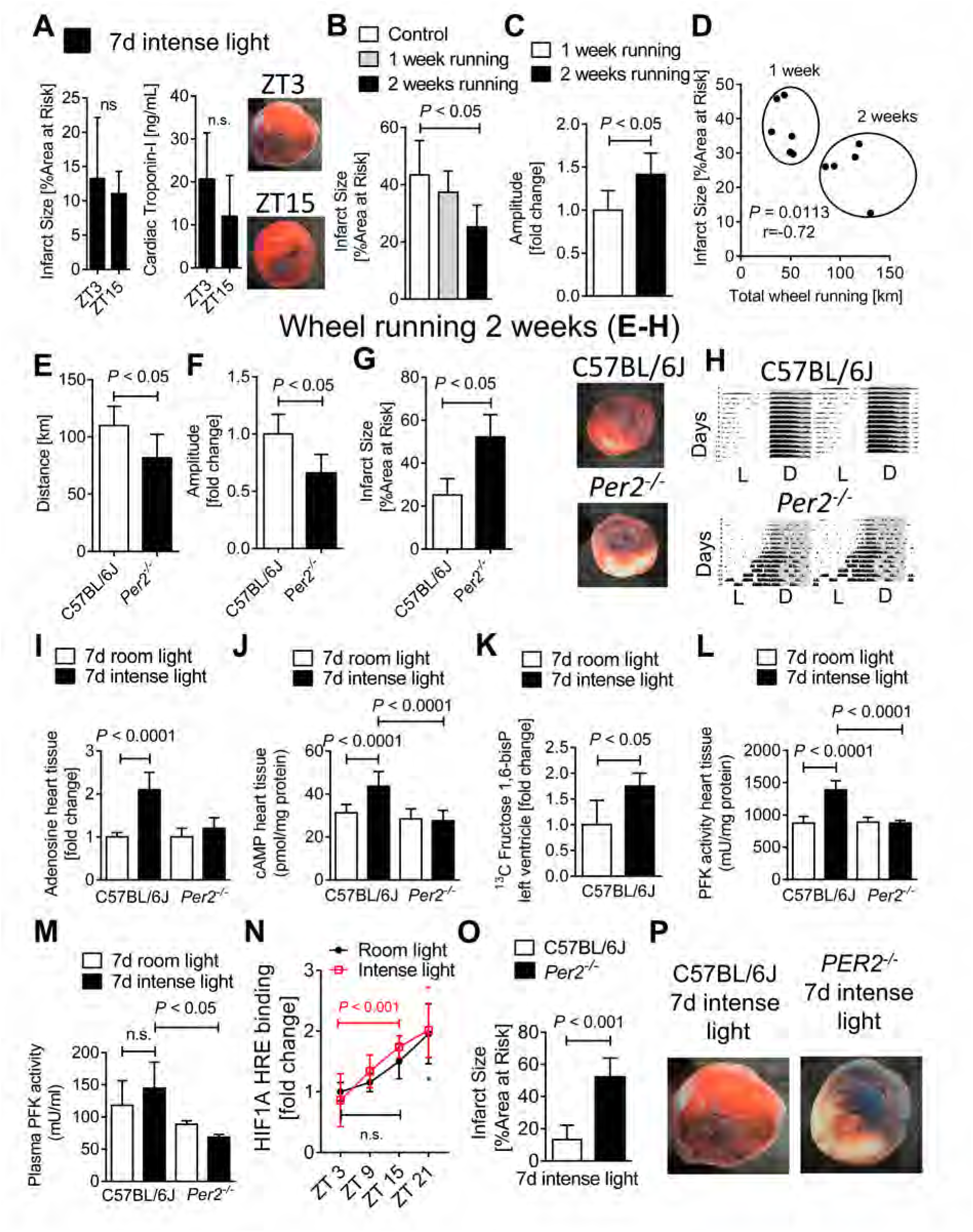
Intense light increases cardiac adenosine-cAMP and glycolytic flux via PER2 in the uninjured heart. **(A)** Infarct sizes in C57BL/6J mice that were housed under intense light (IL, 10,000 LUX, L:D 14:10 h) for 7 days and subjected to 60 min of *in situ* myocardial ischemia followed by 2h reperfusion at ZT3 or ZT15 (mean±SD; n=6; Student’s t test). **(B-D)** C57BL/6J mice exposed to voluntary wheel running for 1 versus 2 weeks. Shown are infarct sizes after 60 min of myocardial ischemia and 2h reperfusion at ZT3 or circadian amplitude and distance walked measurements (mean±SD; n=6; Student’s t test). (**E-H**) Wheel running measurements during or infarct size studies after 2 weeks of wheel running at ZT3 in C57BL/6J or *Per2^-/-^* mice (mean±SD; n=5; Student’s t test). One representative infarct size staining and one wheel running activity recording are shown. **(I, J)** Adenosine or cAMP levels in heart tissue from C57BL/6J or *Per2^-/-^* mice at ZT3 after 7d of room-(RL, 200 LUX, L:D 14:10 h) or intense light (IL, 10,000 LUX, L:D 14:10 h) housing (mean±SD; n=5; ANOVA with Tukey’s multiple comparisons test). **(K)** Cardiac U-^13^C-Fructose-1,6-bisphospahte levels at ZT3 from C57BL/6J mice that were housed under RL or IL for 7 days (mean±SD; n=4; Student’s t test). (**L, M**) Phosphofructokinase (PFK) activity in both heart tissue and plasma samples from C57BL/6J or *Per2^-/-^* mice at ZT3 after 7d of RL or IL housing (mean±SD; n=4-5; ANOVA with Tukey’s multiple comparisons test). **(N)** HIF1A-hyoxia response element (HRE) binding was determined at ZT 3, 9, 15 and 21 (mean±SD; n=5; *Note:* * *P* < 0.05 for ZT21 vs. ZT3 in RL and IL housed mice via Student’s t test. **(O)** C57BL/6J or *Per2^-/-^* mice housed under IL for 7 d prior to 60 min myocardial ischemia and 2 h reperfusion at ZT3; (mean±SD; n=5; Student’s t test). (**P**) Representative infarct staining.

Since intense light had increased the physical activity in mice (Figure 1E), we investigated the influence of voluntary wheel running (Schroeder et al., 2012) on the circadian amplitude and myocardial infarct sizes. In contrast to intense light exposure, however, two weeks of voluntary wheel running with a longer distance walked were necessary until we noted robust cardioprotection from myocardial ischemia (Figure 2B) or a significant increase of the circadian amplitude (Figure 2C). Nevertheless, the total distance achieved on the wheel inversely correlated with infarct sizes (Figure 2D). We, therefore, evaluated two weeks of voluntary wheel running in *Per2^-/-^* mice, which revealed a decrease in total running distance, decreased circadian amplitude, increased infarct sizes and significant circadian disruption in *Per2^-/-^* mice compared to wildtype controls (Figure 2E-H). These findings demonstrate that PER2 is essential for driving the precise rhythmicity of circadian oscillations (Hallows et al., 2013).

As adenosine-mediated increases of cAMP is a core component of PER2 expression and PER2 mediated ischemic preconditioning of the heart (Eckle et al., 2012; O’Neill et al., 2008), we next evaluated if intense light also ‘preconditioned’ the heart similar to ischemia. Analyzing uninjured hearts from wildtype or *Per2^-/-^* mice after one week of intense light ‘preconditioning’ we discovered that intense light significantly increased cardiac adenosine or cAMP levels, which was abolished in *Per2^-/-^* mice (Figure 2I-J). As light elicited adenosine increase was abolished in *Per2^-/-^* mice, and light induction of cardiac PER2 protein required visual perception (Figure 1K), these data suggest intense light elicited adenosine as circulating signaling molecule from the brain (Zhang et al., 2006) to enhance peripheral PER2 expression.

Based on observations that PER2 initiates a switch from energy-efficient lipid to oxygen-efficient glucose metabolism during myocardial ischemia which is pivotal to allow the myocardium to function (Aragones et al., 2009), we next assessed the effect of intense light on glycolytic flux during normoxia. Using liquid chromatography-tandem mass spectrometry studies following the infusion of labeled glucose ([U]^13^C-glucose), we found that intense light significantly increased the glycolytic flux in cardiac tissue before an ischemic event (Figure 2K). We further found that intense light increased the activity of the key and rate-limiting enzyme of glycolysis (phosphofructokinase) in heart tissue or plasma in a PER2 dependent manner (Figure 2L-M).

Considering glycolysis is HIF1A regulated under conditions of low oxygen availability (Krishnan et al., 2009), we next investigated if intense light would increase cardiac HIF1A-hypoxia response element (HRE) binding under normoxia and before an ischemic insult. Indeed, intense light significantly increased total cardiac HIF1A-HRE binding at ZT15 vs. ZT3 when compared to room light conditions (Figure 2N). Finally, subsequent myocardial ischemia and reperfusion studies in *Per2^-/-^* mice confirmed that intense light elicited circadian amplitude enhancement and subsequent cardioprotection was PER2 dependent (Figure 2O, P).

Taken together, these studies found that intense light does not work via increases of physical activity only, but ‘preconditions’ cardiac tissue via increases of cardiac adenosine-cAMP signaling, HIF1A-HRE binding and energy efficient glycolysis prior to an ischemic insult. Furthermore, our data suggest intense light elicited adenosine as circulating signaling molecule (Zhang et al., 2006) to enhance peripheral PER2 mediated cardioprotection.

### Intense light elicited cardioprotection is abolished in mice with endothelial-specific deletion of *Per2*

To further understand intense light elicited and PER2 dependent pathways, we performed a genome-wide array, profiling intense light-dependent gene expression before an ischemic event. *In silico* analysis found dominant regulation of circadian and metabolic pathways (Figure 3A) and identified the hypoxia/HIF1A-regulated and metabolic key player Angiopoietin-Like 4 (ANGPTL4) as the top light and PER2 dependent gene (Figure 3B), supporting our findings that intense light elicited PER2 activates HIF1A regulated pathways under normoxic conditions. As ANGPTL4 is an endothelial secreted protein which protects the endothelial barrier function during myocardial ischemia (Galaup et al., 2012), we next evaluated endothelial specific *Per2* deletion in myocardial IR-injury. Using a tissue-specific mouse line with a 70% deletion of PER2 in the endothelium (Figure 3C, *Per2^loxP/loxP^*-VE-Cadherin-Cre, Figure S2), we found significantly increased infarct sizes and troponin-I serum levels in *Per2^loxP/loxP^*-VE-Cadherin-Cre (Figure 3D-F). In fact, intense light elicited cardioprotection was abolished in *Per2^loxP/loxP^*-VE-Cadherin-Cre mice. As these data implicated intense light in maintaining the vascular integrity during myocardial IR-injury, we determined the vascular leakage of Evans blue dye following 7 days of room light or intense light housing. As shown in Figure 3G-I, intense light ‘pretreatment’ significantly improved endothelial barrier function during myocardial IR-injury which was abolished in endothelial-specific *Per2^-/-^* mice.

**Figure 3.**
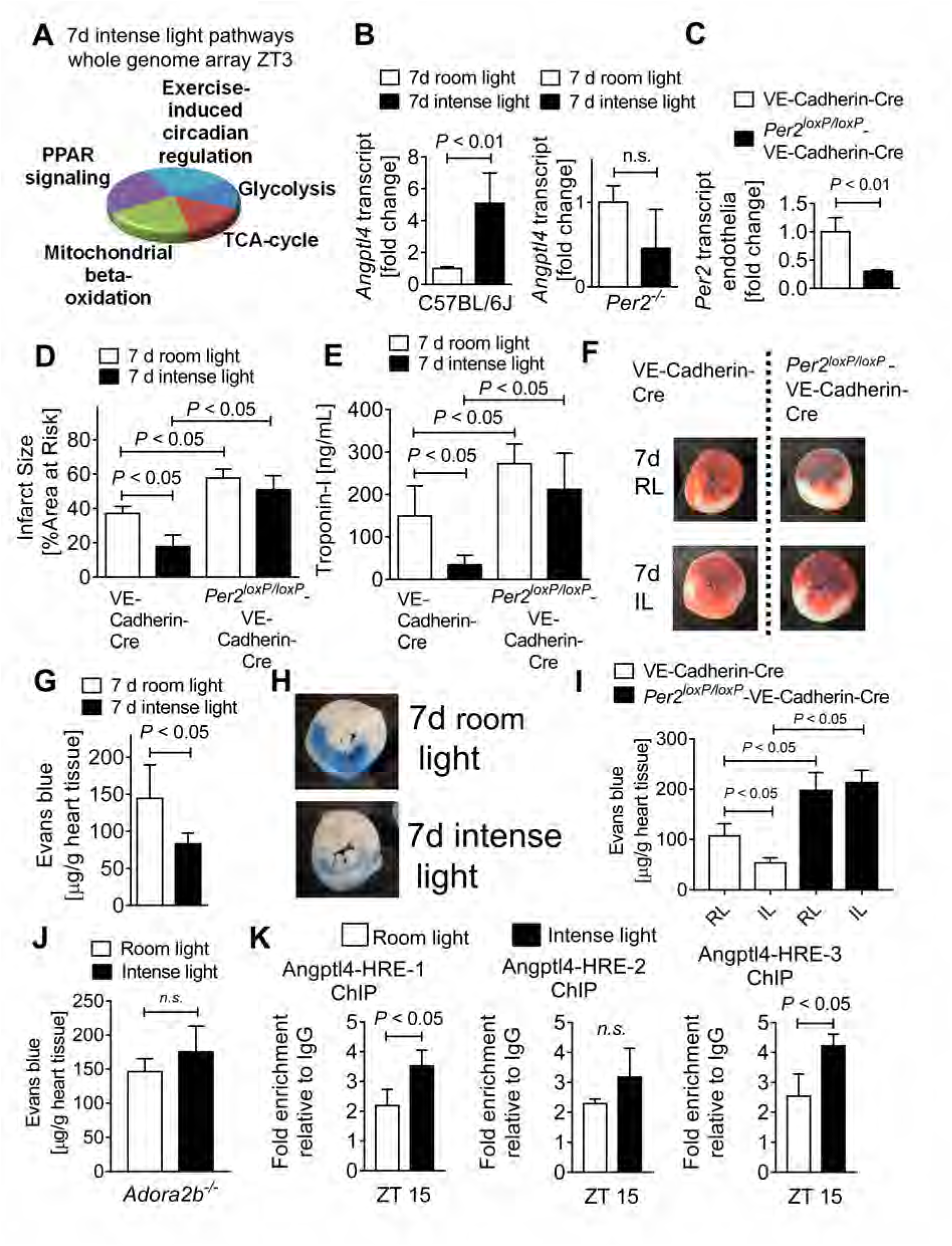
Intense light elicited cardioprotection is endothelial PER2 specific. **(A)** Whole genome array from C57BL/6J or *Per2^-/-^* heart tissue after 7d of intense light (IL, 10,000 LUX, L:D 14:10 h) or standard room light (RL, 200 LUX, L:D 14:10 h) housing at ZT3 (n=3 per group, total of 12 arrays). Shown top light regulated pathways. (**B**) Validation of transcript levels of the top light and PER2 dependent gene (ANGPTL-4) identified by whole genome array (mean±SD; n=4-5; Student’s t test). (**C**) *Per2* mRNA transcript levels from endothelial cells isolated from endothelial specific PER2 deficient (*Per2^loxP/loxP^*-VE-Cadherin-Cre) or control (VE-Cadherin-Cre) hearts (mean±SD; n=3; Student’s t test). (**D, E**) Infarct sizes or serum troponin-I in *Per2^loxP/loxP^*-VE-Cadherin-Cre or VE-Cadherin-Cre mice housed under RL or IL conditions for 7d followed by 60 min of *in situ* myocardial ischemia and 2h reperfusion at ZT3 (mean±SD; n=5; ANOVA with Tukey’s multiple comparisons test). (**F**) Representative infarct staining. (**G-I**) Vascular leakage of Evans blue dye in C57BL/6J or *Per2^loxP/loxP^*-VE-Cadherin-Cre after 60 min of *in situ* myocardial ischemia and 2h reperfusion at ZT3 following 7 days of RL or IL housing (mean±SD; n=5; Student’s t test (**G**) and ANOVA with Tukey’s multiple comparisons test (**I**)). (**J**) Vascular leakage of Evans blue dye in *Adora2b^-/-^* after 60 min of *in situ* myocardial ischemia and 2h reperfusion at ZT3 following 7 days of RL or IL housing (mean±SD; n=5; Student’s t test). (**K**) ChIP assay for HIF1A binding to the promoter region of Angptl4 in C57BL/6J following 7 days of room- RL or IL housing (mean±SD; n=3; Student’s t test). See also Figure S2 and Figure S3.

Recent studies identified adenosine signaling via the adenosine A2B receptor (ADORA2B) as a crucial pathway for PER2 stabilization during myocardial ischemia (Eckle et al., 2012). As intense light had increased cardiac adenosine levels, we questioned if ADORA2B mediated adenosine signaling could be the signaling link between the brain and the heart. Indeed, our data revealed abolished intense light therapy mediated improvement of the endothelial barrier function in *Adora2b^-/-^* mice (Figure 3J).

Considering intense light had increased HIF1A-HRE binding at ZT15, we next evaluated HIF1A binding to the promoter region of mouse *Angptl4* at ZT15. Evaluation of mouse *Angptl4* promoter regions identified several HRE binding sites (Figure S3) and following ChIP assays demonstrated significantly increased HIF1A binding in two promoter regions (Figure 3K).

Taken together these data identify endothelial specific PER2 as a mechanism of intense light elicited cardioprotection and suggest intense light as a strategy to improve endothelial barrier function via increase of adenosine-ADORA2B signaling and HIF1A-transcription.

### Endothelial PER2 is critical for the transcriptional control of HIF1A dependent glycolysis

Based on our findings for a vital role of endothelial-specific PER2 in intense light-mediated cardioprotection and endothelial barrier protection during myocardial IR-injury *in vivo*, we next evaluated endothelial PER2 signaling targets during hypoxia *in vitro*. For this purpose, we generated a lentiviral-mediated PER2 knockdown (KD) stable cell line in human microvascular endothelial cells (HMEC-1). Similar to previous studies in PER2 gene-targeted mice (Eckle et al., 2012), hypoxia increased PER2 transcript or protein levels in HMEC-1 scrambled (Scr) controls, whereas PER2KD HMEC-1 displayed abolished transcriptional induction of HIF1A dependent glycolytic enzymes, attenuated lactate production, reduced glycolytic capacity, and increased cytotoxicity (Figure S4). Based on observations in HMEC-1 that PER2 is significantly increased at 24h after cell synchronization when compared to the 12h time point (Eckle et al., 2012), we determined whether oscillatory higher PER2 levels would affect metabolism under normoxic conditions. These studies revealed a significant increase of glycolytic capacity in control HMEC-1 compared to PER2KD cells at the 24h time point (Figure S4). Mechanistic studies, using a chromatin immunoprecipitation assay (ChIP) uncovered hypoxia induced HIF1A binding to the human lactate dehydrogenase promotor region, a response that was abolished in PER2KD cells (Figure S4).

Together, these findings uncover a critical role for endothelial PER2 in cellular metabolic adaption under normoxia or hypoxia and reveal endothelial PER2 as an essential co-factor of HIF1A mediated transcription of glycolytic genes, and thus a key regulator of glycolytic metabolism.

### Identification of endothelial PER2 as a regulator of TCA cycle activity

As hypoxia increased PER2 protein like intense light, we next used an unbiased affinity purification-mass spectrometry-based proteomics screen for PER2 protein interactions under hypoxic conditions to gain a deeper mechanistic perspective of endothelial PER2 dependent mechanisms (Figure 4A, Table S1, Figure S5). Serendipitously, a high percentage of PER2-protein interactions hinted towards an essential role for PER2 in controlling TCA cycle function (Figure 4B). Subsequent Co-IP pull-downs on TCA cycle enzymes confirmed binding to PER2 during hypoxia (Figure 4C, D). Following analyses of subcellular compartments found that hypoxia increased PER2 protein levels in the cytoplasm, nucleus and the mitochondria (Figure 4E). Thus, PER2 protein interactions may facilitate the transport of mitochondrial proteins, which are almost exclusively synthesized in the cytosol. In fact, our proteomics screen indicated PER2 binding to the mitochondrial outer membrane translocase (Tom) complex [Table S1 (Faou and Hoogenraad, 2012)], which is the main protein entry gate of mitochondria (Boengler et al., 2011). Following co-localization studies confirmed PER2 translocation into the mitochondria during hypoxia (Figure 4F, Figure S6). Functional assays on TCA cycle enzyme activity revealed regulation of TCA cycle function during hypoxia in a PER2 dependent manner (Figure 4G-I), and hypoxic PER2 KD cells showed significantly less CO2 production, a surrogate endpoint of TCA cycle function (Figure 4J).

**Figure 4.**
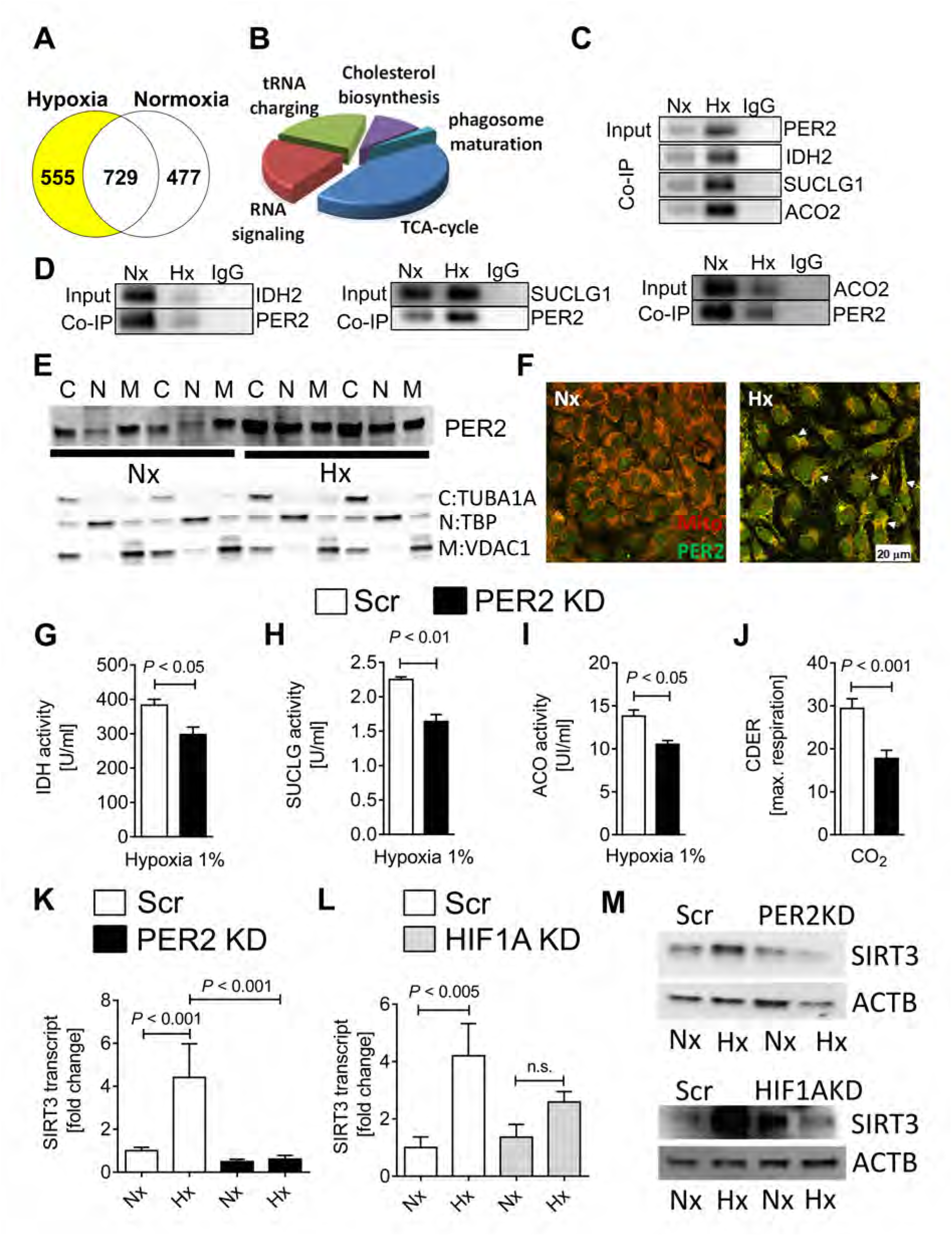
Identification of endothelial PER2 as a regulator of TCA cycle activity. HMEC-1 or stable lentiviral-mediated PER2KD and Scr control HMEC-1 were synchronized and exposed to 24h of normoxia (Nx) or 1% hypoxia (Hx). In a subset of experiments synchronized stable lentiviral-mediated HIF1AKD and Scr HMEC-1 were exposed to Nx or Hx. **(A-B)** Affinity purification-mass spectrometry-based proteomics screen for PER2 protein interactions in normoxic and hypoxic HMEC-1. (**C, D**) Co-immunoprecipitation for PER2 in hypoxic or normoxic HMEC-1 against isocitrate dehydrogenase 2 (IDH2), succinyl Co-A ligase (SUCLG1) and aconitase (ACO2) and vice versa. One representative blot of three is displayed. (**E**) Subcellular compartment analysis of PER2 during normoxia or hypoxia (C= cytoplasm, N=nucleus, M=mitochondria; compartment-specific loading controls: Tubulin Alpha 1a (TUBA1A) for cytoplasm, TATA-Box Binding Protein (TBP) for nucleus and Voltage-Dependent Anion Channel 1 (VDAC1) for mitochondria). (**F**) Translocation of PER2 into the mitochondria during hypoxia. **(G-I)** TCA cycle enzyme activities of IDH, SUCLG and ACO from stable lentiviral-mediated PER2KD and Scr control HMEC-1 during hypoxia (mean±SD, n=3; Student’s t test). **(J)** Carbon dioxide evolution rates (CDER), as a surrogate for TCA cycle function, in PER2 KD or Scr HMEC-1 measured by a mitochondrial stress test using a Seahorse XF24 FluxPak assay (mean±SD, n=5; Student’s t test). (**K-M**) SIRT3 transcript or protein levels from stable lentiviral-mediated PER2KD and Scr or stable lentiviral-mediated HIF1AKD and Scr control HMEC-1 (mean±SD, n=3; ANOVA with Tukey’s multiple comparisons test). See also Figure S4, Figure S5, Figure S5 and Figure S7.

Considering TCA cycle enzyme activity is also known to be regulated by Sirtuin3 (SIRT3) mediated de-acetylation (Yu et al., 2012), which is under circadian control (Peek et al., 2013), we investigated whether hypoxia and HIF1A-PER2 dependent pathways would regulate SIRT3 expression. HMEC-1 transcriptional or translational analyses with a PER2- or HIF1A KD revealed a PER2-HIF1A dependent regulation of SIRT3 under hypoxic conditions (Figure 4K-M, Figure S7). *In silico* analysis confirmed a hypoxia response element (HRE) in the human promoter region of SIRT3 (Figure S7).

Together, our proteomics screen uncovered a critical role for endothelial PER2 in controlling oxidative TCA cycle metabolism during hypoxia by translocating into the mitochondria and via transcriptional regulation of HIF1A-SIRT3-dependent pathways. These data suggest a more complex function of PER2, possibly controlling the TCA cycle function via post-translational mechanisms.

### Endothelial PER2 transcriptionally regulates mitochondrial respiration and barrier function

Additional analysis of our proteomics screen indicated binding of PER2 to mitochondrial complex 4 (Table S1, *Cytochrome C*), supporting a role for PER2 in controlling mitochondrial function under hypoxia. Indeed, oxygen consumption rates (OCR; a measure of mitochondrial functionality), basal respiration, maximal respiration, ATP production, and spare capacity were significantly reduced in PER2KD cells during a mitochondrial stress test (Figure 5A-D, Figure S7). Moreover, OCR levels were significantly increased in cells with higher PER2 levels at time point 24h when compared to 12h post cell synchronization (Eckle et al., 2012). These findings highlight a role for oscillatory PER2 overexpression in metabolic adaptation under normoxia (Figure S7).

**Figure 5.**
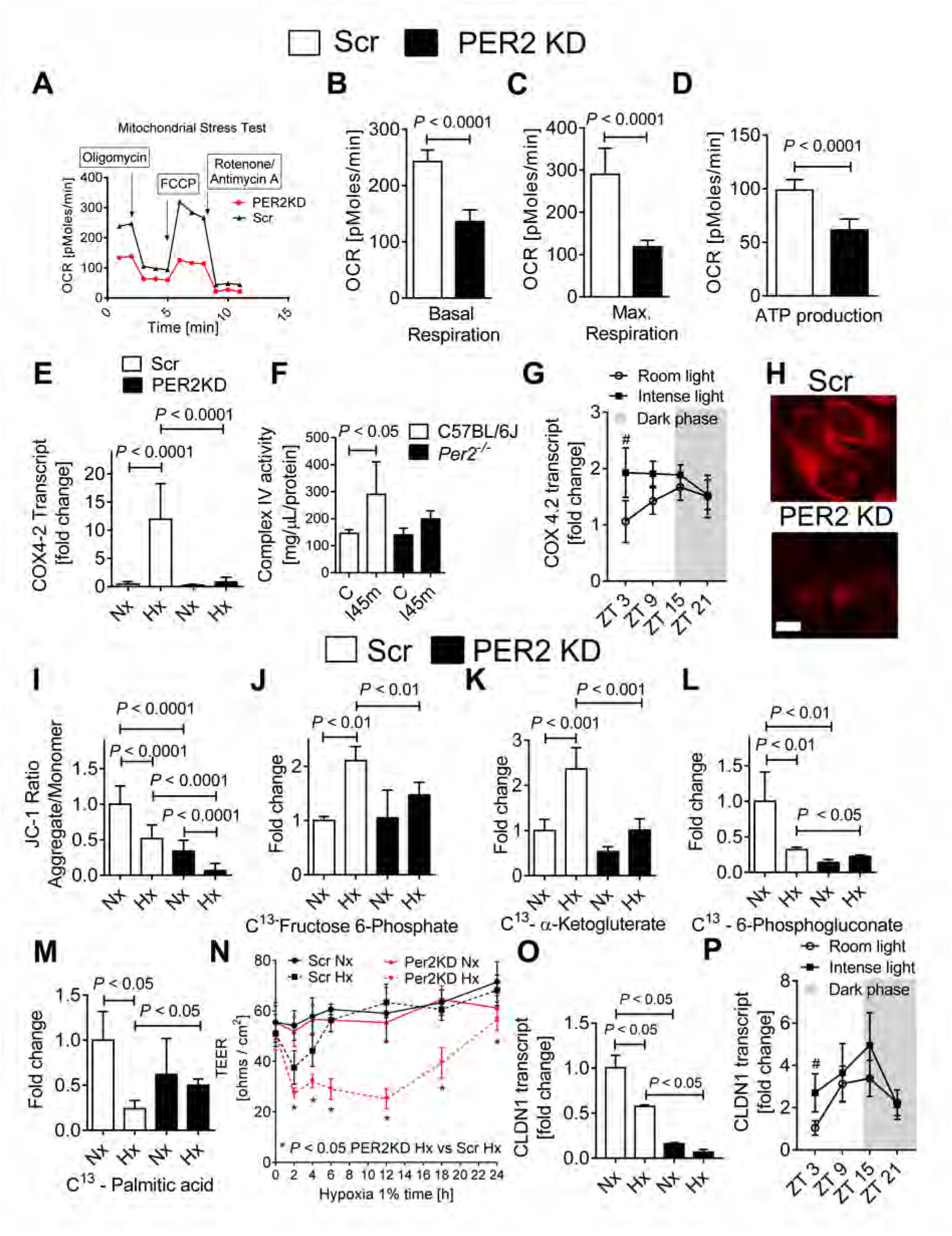
Endothelial PER2 regulates mitochondrial ATP production and barrier function. **(A-D)** Oxygen consumption rates (OCR) in PER2KD or Scr HMEC-1. Quantification of basal respiration, maximum achievable respiration, and ATP production are shown (mean±SD, n=5; Student’s t test). **(E**) COX4.2 transcript levels in PER2 KD or Scr HMEC-1 after 24h of Nx or 1% Hx treatment (mean±SD, n=6; ANOVA with Tukey’s multiple comparisons test). **(F)** Complex IV enzyme activity in *Per2^-/-^* or C57BL/6 mouse hearts subjected to 45 min of ischemia (mean±SD, n=4; ANOVA with Tukey’s multiple comparisons test). **(G)** Cardiac *Cox42 mRNA* levels at ZT 3, 9, 15 and 21 in C57BL/6 mice after 7d or room light (RL) or intense light (IL) housing (mean±SD; n=5) *Note:* ^#^ *P* < 0.05 for ZT3 IL vs. ZT3 in RL housed mice via Student’s t test. **(H)** MitoTracker Red CMXRos staining of PER2 KD or Scr HMEC-1 at baseline. One representative image of 5 is shown. (**I**) Quantification of the mitochondrial membrane potential probe JC-1 (mean±SD, n=6; ANOVA with Tukey’s multiple comparisons test). **(J-M)** ^13^C-metabolites from supernatants of PER2 KD or Scr HMEC-1 following 24h of Nx or 1% Hx treatment. Data are presented as percentage of total metabolite present (mean±SD, n=3; ANOVA with Tukey’s multiple comparisons test). (**N**) Permeability assay in PER2 KD or Scr HMEC-1 during 24h of 1% hypoxia (mean±SD, n=5; ANOVA with Tukey’s multiple comparisons test). *Note:* Permeability increases after prolonged hypoxia exposure of endothelial cells due to morphological changes (Am. J. Physiol. 269 (Lung Cell. Mol. Physiol. 13): L52-L58, 1995.). (**O**) *CLDN1* (Claudin-1) transcript levels in PER2 KD or Scr HMEC-1 after 4h of Nx or 1% Hx treatment (mean±SD, n=3; ANOVA with Tukey’s multiple comparisons test). (**P**) Cardiac *Cldn1 mRNA* was determined at ZT 3, 9, 15 and 21 in C57BL/6 mice after 7d or RL or IL treatment (mean±SD; n=5; *Note:* ^#^ *P* < 0.05 for ZT3 IL vs. ZT3 in RL housed mice via Student’s t test). See also Figure S8 and Figure S9.

Considering HIF1A mediates a switch of complex 4 subunits (COX4.1 to COX4.2) in hypoxia to enhance oxygen efficiency, which conserves cellular ATP content (Fukuda et al., 2007), we next investigated the transcriptional regulation of COX4.2 in PER2KD cells under hypoxia. Here, we found abolished increases of COX4.2 mRNA or complex 4 activity in hypoxic PER2KD cells or ischemic hearts from *Per2^-/^*^-^ mice, respectively (Figure 5E, F). Moreover, intense light ‘preconditioning’ of wildtype mice resulted in significantly increased cardiac COX4.2 mRNA levels at ZT3 in the uninjured heart (Figure 5G).

To understand if compromised oxidative phosphorylation in PER2 deficiency would be associated with a reduced mitochondrial membrane potential, which is associated with compromised mitochondrial function (Solaini et al., 2010), we next used MitoTracker deep red staining (Zhou et al., 2011). Studies in PER2KD HMEC-1 indicated already reduced mitochondrial potential under normoxia (Figure 5H). Indeed, analysis of a cell energy phenotype assay revealed significantly less aerobic metabolism in PER2KD cells at baseline (Figure S8). Confirming these results, JC-1 assay showed a significant reduction of the membrane potential in PER2 KD cells at both normoxia and under hypoxia (Figure 5I, Figure S8).

To further explore PER2 dependent metabolism, we next used liquid chromatography-tandem mass spectrometry studies following the exposure of labeled glucose (^13^C-glucose) or palmitic acid (^13^C-palmitic acid) to assess metabolic flux in PER2KD endothelial cells. Here we confirmed that PER2 is an essential regulator of glycolysis and oxidative metabolism under hypoxia (Figure 5J-K). Moreover, we also found PER2 to be critical for the pentose phosphate pathway under normoxia or hypoxia, indicating that PER2KD cells are compromised in generating the redox cofactor NADPH, which has a pivotal role for circadian timekeeping (Figure 5L) (Rey et al., 2016). As PER2 has been shown to inhibit lipid metabolism via PPARγ (Grimaldi et al., 2010), we also found altered fatty acid metabolism in PER2KD cells under hypoxia (Figure 5M). *In silico* analysis of our proteomics screen confirmed these findings and highlight PER2 as a master regulator of endothelial energy metabolism (Figure S9).

As ATP has been implicated in endothelial barrier enhancement and tight junction functionality (Kolosova et al., 2005), we next evaluated endothelial barrier function of PER2KD HMECs and controls during a 24h hypoxia time course. As shown in Figure 5N, PER2KD HMEC demonstrated increased cell permeability at 2, 4, 6, 12 and 24 h of hypoxia when compared to scrambled controls. Considering previous studies had demonstrated that HIF-dependent regulation of claudin-1 is central to epithelial tight junction integrity (Saeedi et al., 2015), we also evaluated claudin-1 expression levels. As shown in Figure 5O, PER2 KD cells had significantly less HIF1A-regulated *Claudin-1* mRNA levels. In addition, intense light ‘preconditioning’ of wildtype mice resulted in significantly increased cardiac *Claudin-1* mRNA levels at ZT3 (Figure 5P).

Taken together, these data identify endothelial PER2 as critical control point of energy homeostasis and endothelial barrier function via transcriptional regulation of HIF1A dependent mitochondrial respiration and claudin-1.

### A light-sensing human endothelial cell line recapitulates *in vivo* light exposure

As proof of concept that PER2 mimics HIF1A pathways under normoxia, we reiterated light sensing for PER2 overexpression on a cellular level by generating an HMEC-1 line, overexpressing the human light sensing photopigment melanopsin (OPN4), a retinal ganglion cell receptor responsible for circadian entrainment. Exposing the light sensing HMEC-1 cultures to light resulted in a significant increase of cAMP, phospho-CREB (cyclic AMP-responsive element binding protein), PER2 mRNA, glycolytic capacity and oxygen consumption rates (Fig. 6A-H). Taken together, these studies recapitulate that normoxic PER2 overexpression can optimize cellular metabolism in a similar fashion as seen under hypoxic conditions.

**Figure 6.**
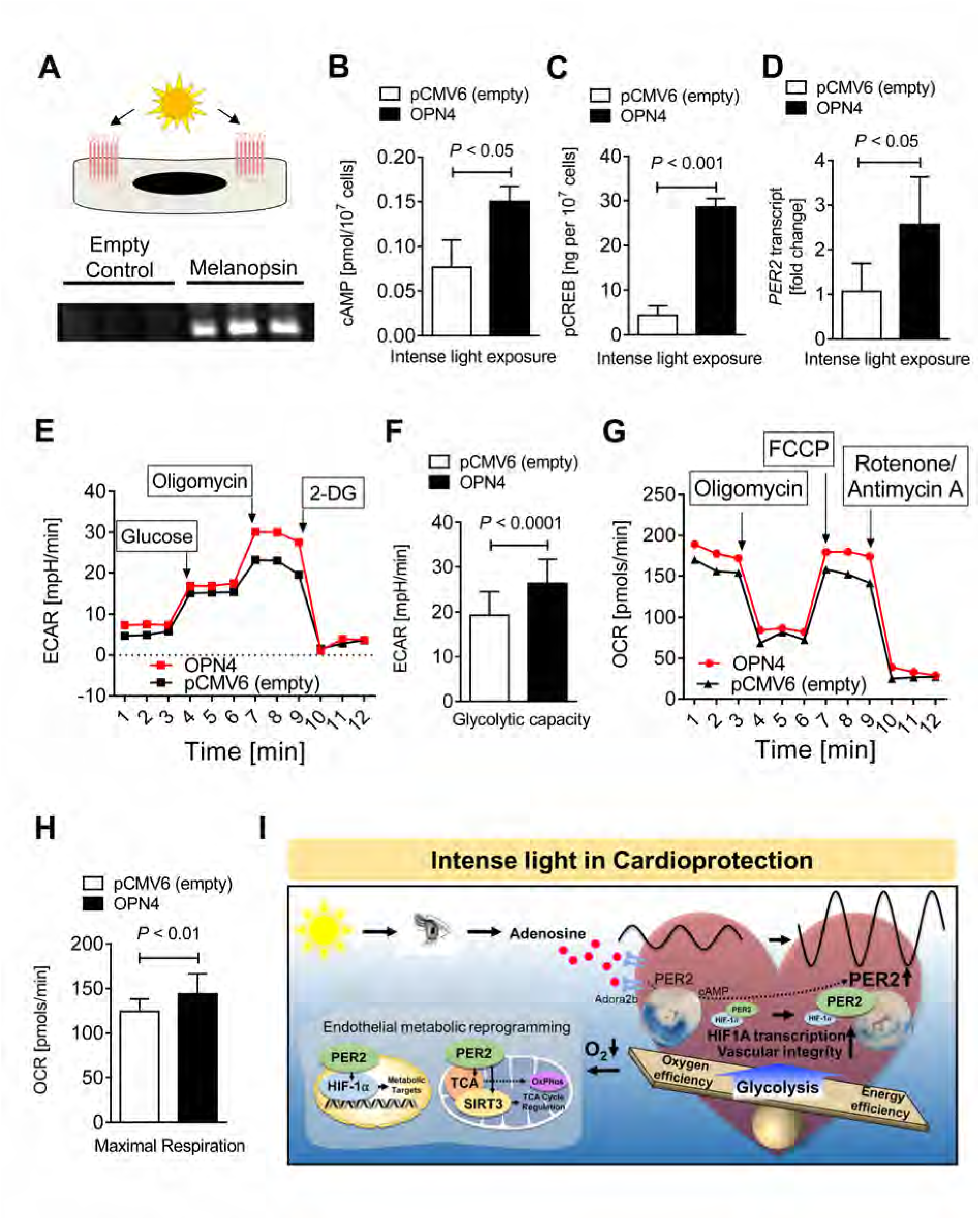
Light-sensing human cell line recapitulates *in vivo* light exposure. (**A**) Study design and verification of melanopsin overexpression by immunoblot. pCMV6 is the empty vector control, and OPN4-pCMV6 is the plasmid containing the gene encoding melanopsin (n=3). (**B-H**) cAMP, pCREB levels, PER2 transcript, glycolytic capacity, and maximum achievable respiration after light-sensing cells were exposed to intense light (mean±SD, n=6-10; Student’s t test). (**I**) Schematic model.

In summary, our *in vivo* and *in vitro* studies on light-elicited pathways identified a light perception dependent circadian entrainment mechanism through adenosine-cAMP and HIF1A transcriptional adaptation in a PER2 regulated manner. Furthermore, our studies discover that light or hypoxia elicits PER2 as a critical factor in maintaining endothelial barrier function during myocardial ischemia via transcriptional reprogramming (Fig. 6I).

### Intense light enhances the circadian amplitude and PER2 dependent metabolism in humans

Next, we investigated if intense light would have similar effects on healthy human volunteers. Based on strategies using intense light therapy [10,000 LUX] to treat seasonal mood disorders in humans (Yorguner Kupeli et al., 2017), we adopted a similar protocol. We exposed healthy human volunteers to 30 min of intense light in the morning on 5 consecutive days and performed serial blood draws. Intense light therapy increased PER2 protein levels in human buccal or plasma samples in the morning (9AM) or evening (9PM), indicating an enhancement of the circadian amplitude in different tissues at the same time via light therapy (Figure 7A-C, Figure S10). To test the efficacy of intense light therapy on the circadian system (Lewy et al., 1980), we determined melatonin plasma levels, which were significantly suppressed upon light treatment (Figure 7D, E). Also, room light was less efficient than intense light therapy in suppressing melatonin (Figure 7D).

**Figure 7.**
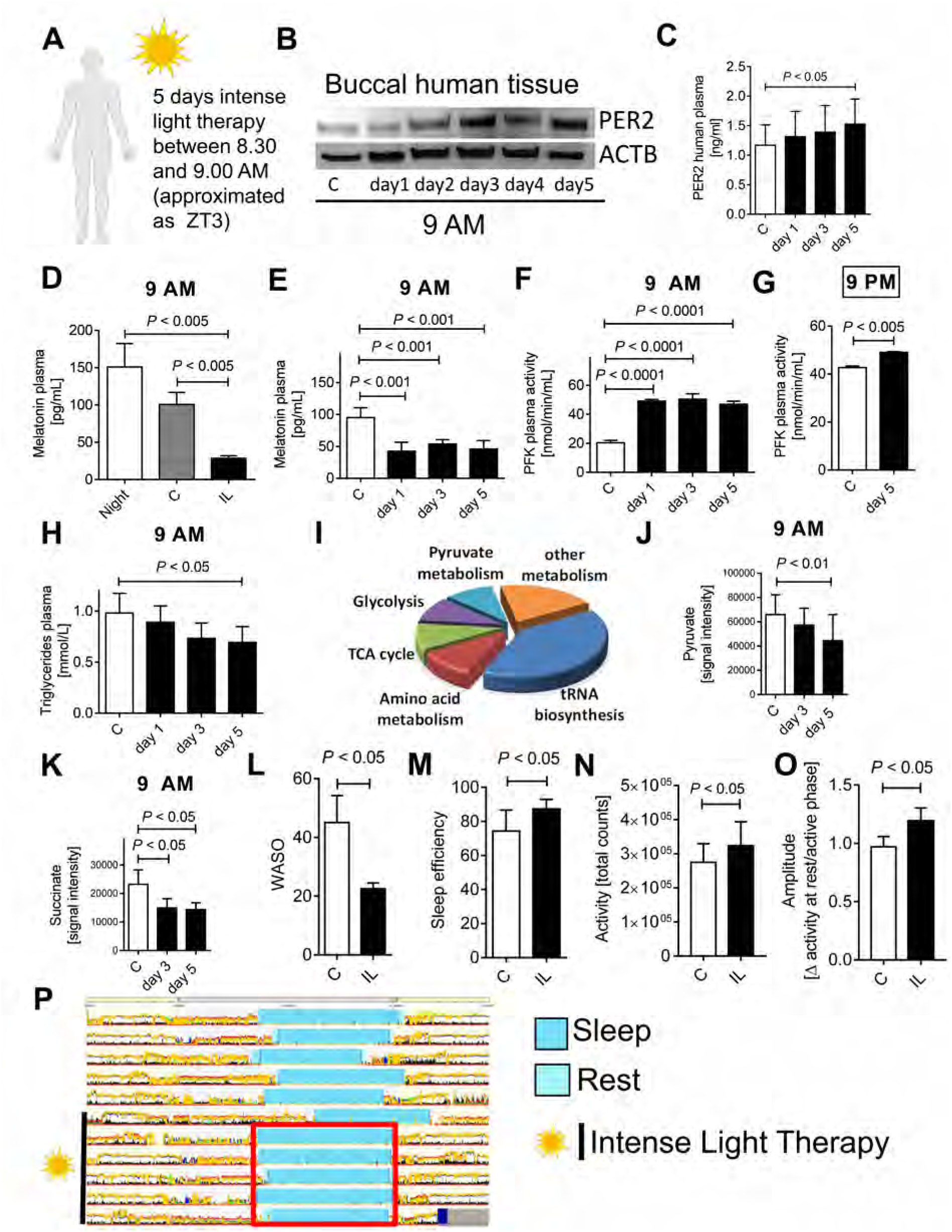
Intense light enhances the circadian amplitude of PER2 and activates PER2 metabolism in humans. **(A)** Protocol for intense light exposure experiments in healthy human volunteers. 20 healthy volunteers (11 female and 6 male, age range between 21-44 yrs.) were exposed to intense light (10,000 LUX) from 8.30-9.00 AM on 5 consecutive days. **(B, C**) PER2 protein levels from buccal tissue or plasma samples at 9AM during 5d of intense light exposure assessed by immunoblot or ELISA, respectively (mean±SD; n=6; ANOVA with Tukey’s multiple comparisons test). **(D)** Effect of room light versus intense light on human plasma melatonin levels (mean±SD; n=3-6; ANOVA with Tukey’s multiple comparisons test). (**E**) Longitudinal monitoring of human plasma melatonin levels during 5d of intense light exposure at 9AM (mean±SD; n=3-6; ANOVA with Tukey’s multiple comparisons test). **(F)** Human plasma phosphofructokinase (PFK) activity during 5d of intense light exposure at 9AM (mean±SD; n=3-6; ANOVA with Tukey’s multiple comparisons test). (**G**) Human plasma PFK activity after 5d of intense light exposure at 9PM (mean±SD; n=3; Student’s t test). **(H)** Human plasma triglyceride levels during 5 d of intense light exposure at 9 AM (mean±SD; n=8; ANOVA with Tukey’s multiple comparisons test). **(I-K)** Targeted metabolomics using mass-spectrometry on human plasma samples from healthy volunteers exposed to intense light therapy for 5 days. Key metabolites of glycolysis (pyruvate) or TCA cycle (succinate) are shown for day 3 and day 5 of intense light therapy (mean±SD; n=3; ANOVA with Tukey’s multiple comparisons test). **(L-P)** Actigraphy data using a validated accelerometer (Actiwatch 2). Shown are the wake-up episodes after the sleep onset (WASO, **L**), the sleep efficiency (**M**), day activity (**N**), the circadian amplitude (**O**, mean±SD; n=6; Student’s t test), and one representative actigraphy recording from one healthy volunteer before and during intense light therapy (**Note**: synchronized sleep phases [turquoise bar] during intense light exposure [red square]). C=control subjects prior to any light exposure, IL=intense light. See also Figure S10 and Figure S11.

Further analyses revealed that intense light therapy increased plasma phosphofructokinase at 9AM or 9PM (Figure 7F, G). Moreover, plasma triglycerides, a surrogate for insulin sensitivity and carbohydrate metabolism (Ginsberg et al., 2005), significantly decreased upon light therapy (Figure 7H), indicating increased insulin sensitivity and glucose metabolism. Targeted metabolomics from human plasma samples confirmed a strong effect of light therapy on metabolic pathways such as glycolysis or TCA cycle (Figure 7I, Figure S10). We found significant decreases in pyruvate or succinate levels after 5 days of light therapy (Figure 7J-K). Together with increased plasma phosphofructokinase activity, this finding indicates improved metabolic flux, possibly due to increased glycolysis, improved TCA cycle or mitochondrial function.

As sleep deprivation is directly associated with decreased insulin sensitivity and compromised glucose metabolism (Depner et al., 2014), we next determined how light therapy would impact human physiology in terms of sleep behavior. Using a validated accelerometer for actigraphy (Lee and Suen, 2017) (Actiwatch 2) we found less WASO (wake after sleep onset) episodes, overall improved sleep efficiency, increased day-activity and increases of the circadian amplitude (Figure 7L-P, Figure S11).

Taken together, our data suggest that intense light therapy, a mechanism of circadian amplitude enhancement, targets similar PER2 dependent metabolic pathways in humans as seen in mice and may present a promising strategy for the treatment or prevention of low oxygen conditions such as myocardial ischemia.

## DISCUSSION

Our studies established a critical role for intense light in regulating critical biological processes (Zadeh et al., 2014). Epidemiologic studies noting an increase in MIs during the darker winter months in *all* US states (Spencer et al., 1998) further support our conclusion that intense light elicits robust cardioprotection. The mechanism of how the entrainment signal gets to peripheral organs remains unclear but may incorporate neuro-hormonal factors, or autonomic innervation (Takahashi, 2017). Studies on altered liver metabolism in constant darkness found adenosine as a possible circulating circadian factor (Zhang et al., 2006), which suggests adenosine signaling as a mechanism for establishing circadian rhythmicity between peripheral organs and the suprachiasmatic nuclei (SCN). Indeed, the importance of adenosine signaling via the adenosine A2B receptor (ADORA2B) for cAMP increases, PER2 stabilization and cardiac metabolic adaption to ischemia has been shown in recent studies investigating the mechanism of myocardial ischemic preconditioning (Eckle et al., 2012). In the current studies, we found that light increased cardiac adenosine and cAMP levels under normoxia, which was also PER2 dependent. While we did not determine plasma adenosine levels, we were able to detect adenosine increases in blood containing and flash frozen mouse hearts. Since intense light pre-treatment did not improve the endothelial barrier function in *Adora2b* deficient mice, adenosine signaling might play an essential role in transmitting the ‘cardioprotective’ light signal from the SCN to the heart. However, as our studies are limited to observations in wildtype, whole body *Adora2b^-/-^* or *Per2^-/-^* and endothelial specific *Per2* deficient mice, future studies will be necessary to investigate the brain specific role of adenosine signaling for peripheral PER2 stabilization.

While cardiomyocytes are significant oxygen consumers and account for approximately 75% of the myocardial volume, there is at least one capillary adjacent to every cardiomyocyte, and cardiomyocytes are outnumbered 3:1 by the endothelial cells (Brutsaert, 2003). Indeed, mitochondrial metabolism in endothelial cells has been proposed as a central oxygen sensor in the vasculature (Davidson and Duchen, 2007), and studies have suggested that human endothelial cells can regulate the activity of HIF1A, thus affecting key response pathways to hypoxia and metabolic stress (Davidson and Duchen, 2007). As such, endothelial dysfunction plays a significant role in myocardial IR-injury, rendering endothelial cells an attractive target for myocardial protection (Yang et al., 2016). In the current studies, we uncovered a critical role for light elicited PER2 in controlling endothelial barrier function. While PER2 has been implicated in endothelial function in previous studies (Viswambharan et al., 2007; Wang et al., 2008), an endothelial specific role of PER2 during acute myocardial IR-injury, which can be targeted using intense light, has not yet been described. Moreover, our *in vivo* and *in vitro* studies suggest that light elicited transcriptional reprogramming of endothelial cells protects the endothelial barrier function during IR-injury. Together with previous studies on myocardial IR-injury (Yang et al., 2016), these studies highlight the importance of cardiac endothelia in IR-injury and point towards unrecognized therapeutic strategies for cardiovascular disease using intense light or pharmacological compounds, such as the circadian rhythm enhancer nobiletin (Gile et al., 2018; Oyama et al., 2018), to increase the amplitude of endothelial PER2. Despite this strong evidence, our studies are limited to the analysis of endothelia specific PER2 deficient mice and thus the contribution of other cardiac cells cannot be fully excluded (Seo et al., 2015).

The importance of HIF1A in cardioprotection has been shown in numerous studies (Semenza, 2014) and the interaction between HIF1A and PER2 has been demonstrated on the protein (Eckle et al., 2012; Kobayashi et al., 2017) and transcriptome level (Wu et al., 2017), In general, HIF1A requires hypoxic conditions to be stabilized (Semenza, 2014). In the current studies, we further established the dependence of HIF1A on PER2 as a transcription factor during hypoxia, supporting previous studies on PER2 function as an effector molecule for the recruitment of HIF1A to promoter regions of its downstream genes (Kobayashi et al., 2017).

However, we also found that specific HIF1A pathways that control glycolysis, mitochondrial respiration (COX4.2) or endothelial barrier function (ANGPTL4/CLDN1) can be transcriptionally activated via light elicited circadian overexpression of PER2 under normoxia. These findings would suggest that PER2 amplitude enhancement strategies can ‘precondition’ the myocardium by establishing a HIF1A-similar signaling environment before an ischemic event. While the role of CLDN1 or COX4.2 in cardioprotection has not been investigated yet, studies have shown the importance of endothelial expressed HIF1A in ischemic preconditioning of the heart (Sarkar et al., 2012). Limitations of these findings include that we did not investigate these pathways in endothelial specific *Per2* or *Hifa* deficient mice. Furthermore, the mechanism how light or hypoxia facilitate a PER2-HIF1A interaction remains elusive. Future genetic studies using tissue specific mice or CRISPR technology to manipulate specific PER2 or HIF1A sequences (Schmutz et al., 2010) will be necessary to understand the role of these pathways and its mechanisms in intense light elicited cardioprotection.

Given a close association of circadian amplitude dampening and disease progression (Gloston et al., 2017), ‘clock’-enhancing strategies are promising approaches for disease treatment. Although it is well known that light regulates circadian rhythms (Czeisler et al., 1990) and that high intensities of light are more effective for circadian entrainment and amplitude enhancement (Lewy et al., 1980), only a few reports exist on circadian entrainment and cardioprotection (Martino et al., 2007). While previous studies suggested that short term intense light exposure could mediate cardioprotection in a PER2 dependent manner (Eckle et al., 2012), no specific mechanisms were provided. In the current studies, we uncovered that intense light increases the circadian amplitude in a PER2 dependent manner, which appeared to be more efficient than exercise induced amplitude enhancement. This finding could have implications for current clinical practice where exercise limitations or the lack of motivation for exercise are commonly observed. However, as we did not investigate blindness in endothelial specific PER2 deficient mice or evaluate different time points for PER2 expression in our blind mice, future studies will be necessary to fully elucidate the role of circadian PER2 amplitude enhancement in cardioprotection.

Interestingly, our light exposure strategy in humans showed similar kinetics as in mice, which could be explained by the fact that circadian rhythms function independently of a diurnal or nocturnal behavior due to multiple yet parallel outputs from the SCN (Kronfeld-Schor et al., 2013). Indeed, very basic features of the circadian system are the same in apparently diurnal and nocturnal animals, including the molecular oscillatory machinery and the mechanisms responsible for pacemaker entrainment by light (Kronfeld-Schor et al., 2013). Also, PER2 is hypoxia-regulated in mice and humans, which supports similar mechanisms in both species (Eckle et al., 2012). HIF1A regulation and function under hypoxia, which is strongly associated with PER2 (Kobayashi et al., 2017), also seems to be independent of a nocturnal nature (Semenza, 2014), despite HIF1A expression being under circadian control (Wu et al., 2017). Indeed, human and mouse studies on HIF1A find similar responses to cardiovascular ischemic events (Semenza, 2014). Nevertheless, differences in size and physiology, as well as variations in the homology of targets between mice and humans, may lead to translational limitations.

Supporting the importance of circadian rhythms in myocardial susceptibility to ischemia, recent studies found a diurnal pattern for troponin values in patients undergoing aortic valve replacement (Montaigne et al., 2018). Here, troponin values following surgery were significantly higher in the morning when compared to the afternoon. While nothing can be done about a diurnal pattern, applying light therapy before high risk non-cardiac or cardiac surgery to enhance the circadian amplitude, however, might be able to provide robust cardioprotection. Light elicited circadian amplitude enhancement suggests an overall increase in PER2 levels and concomitant cardioprotection even at the trough of the amplitude, indicating that this strategy could promote general cardioprotection and potentially decrease troponin levels in both the morning and evening times. However, future studies in humans will be necessary to understand the impact of intense light therapy and its potential role in cardioprotection.

## Acknowledgement

The authors wish to acknowledge Melissa Card, the University of Colorado Molecular and Cellular Analytical Core of the Colorado Nutrition and Obesity Research Center for use of the Seahorse Bioanalyzer, and the University of Colorado School of Medicine Biological Mass Spectrometry Core Facility for technical assistance.

## Author contributions

Y.O.: wrote the manuscript, performed experiments, analyzed the data; C.M.B: wrote the manuscript, performed experiments, analyzed the data; S.B.: wrote the manuscript, performed experiments, analyzed the data; J.S.L. : wrote the manuscript, performed experiments, analyzed the data; L.A.W. wrote the manuscript, performed experiments, analyzed the data; J.H. wrote the manuscript, performed experiments, analyzed the data; C.H.B wrote the manuscript, performed experiments, analyzed the data; P.M.B.: wrote the manuscript; C.M.A.: wrote the manuscript; N.C.: wrote the manuscript, performed experiments, analyzed the data; S.P.C: wrote the manuscript; T.E.: designed the study, wrote the manuscript, performed experiments, analyzed the data.

## STAR METHODS

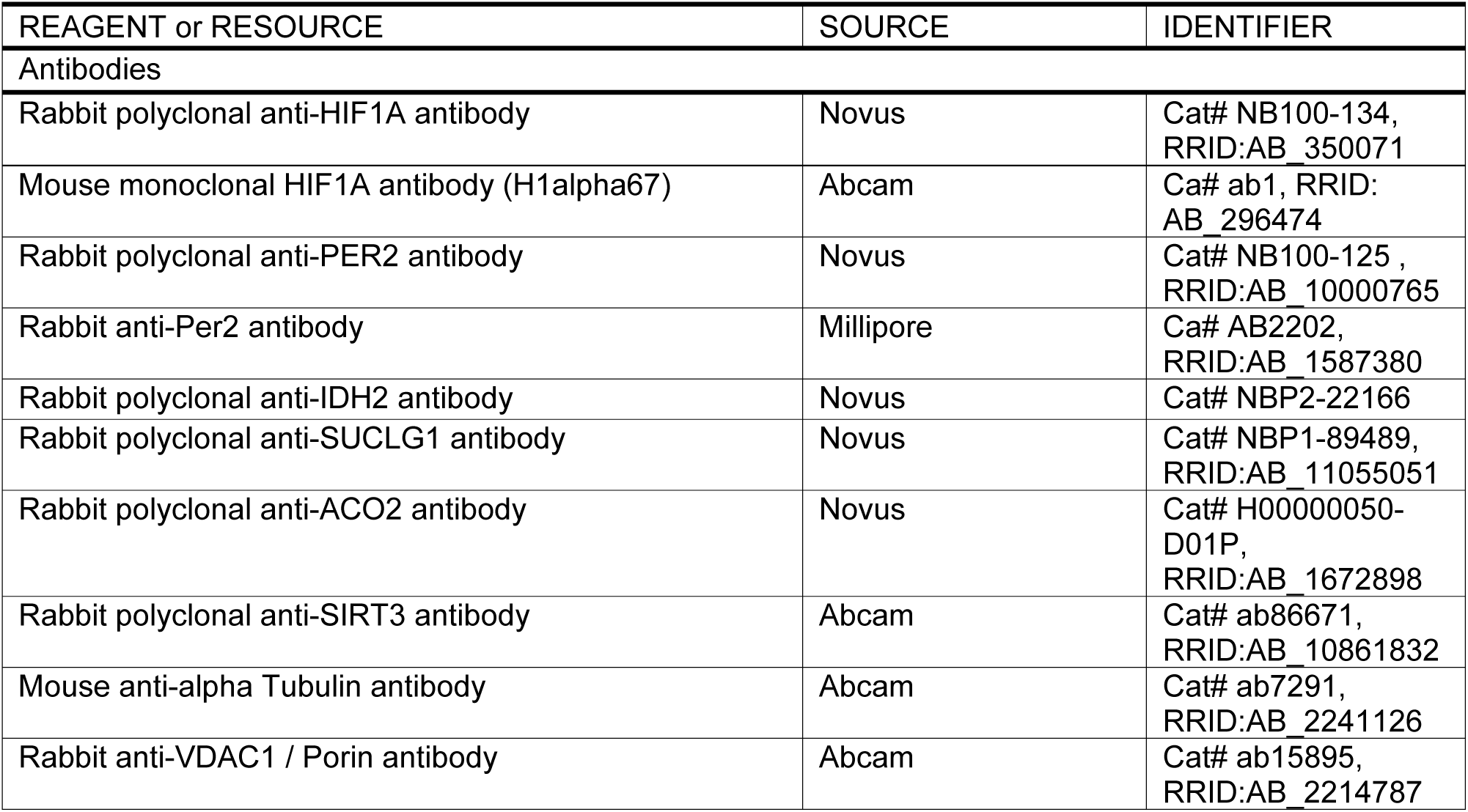
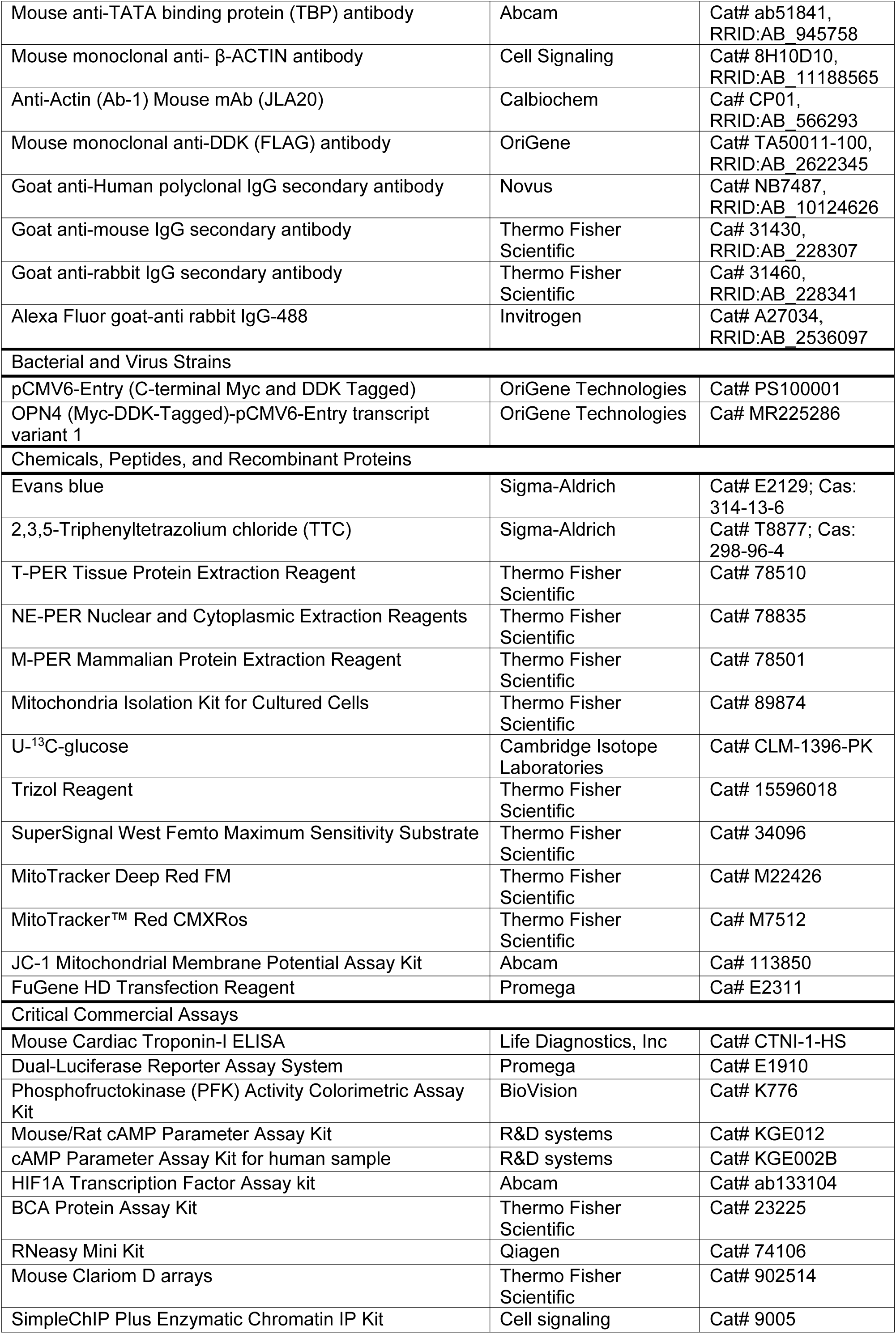
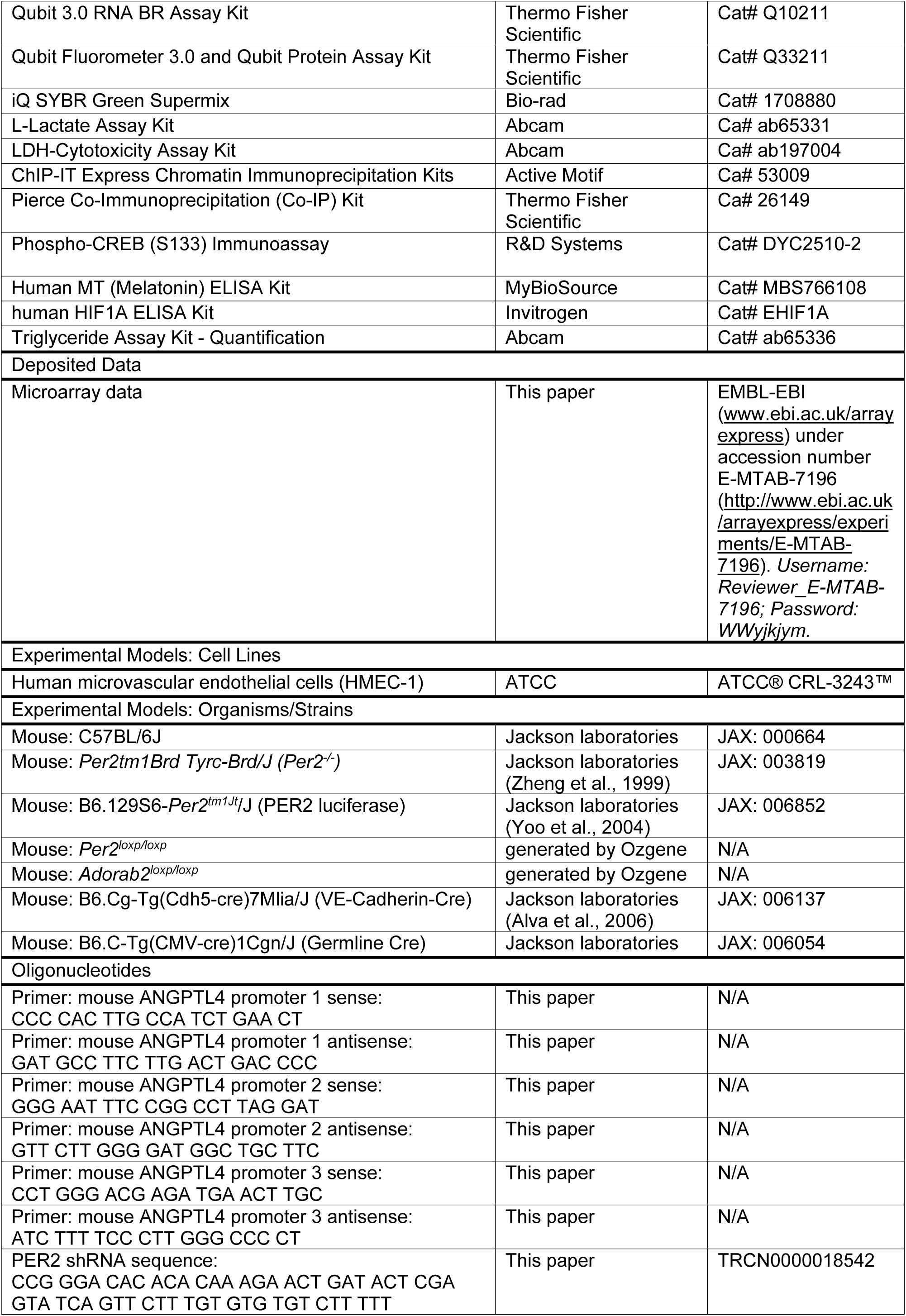
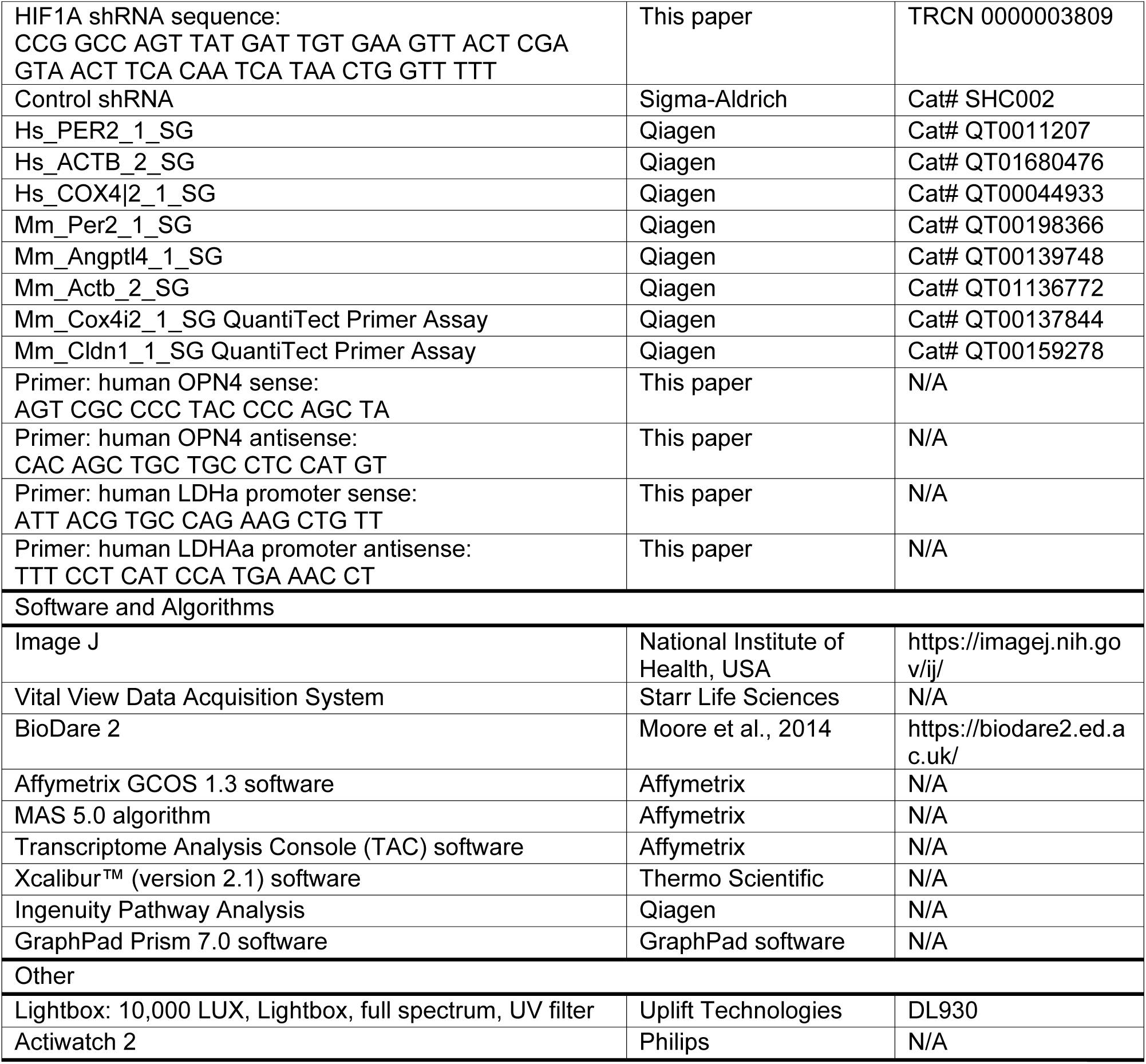
KEY RESOURCE TABLE.

## LEAD CONTACT AND MATERIALS AVAILABILITY

Further information and requests for resources should be directed to and will be fulfilled by the Lead Contact, Tobias Eckle. Mouse and cell lines generated in this study are available upon request via a material transfer agreements (MTA).

## EXPERIMENTAL MODEL AND SUBJECT DETAILS

### Mouse experiments

Experimental protocols were approved by the Institutional Review Board (Institutional Animal Care and Use Committee [IACUC]) at the University of Colorado Denver, USA. They were in accordance with the NIH guidelines for the use of live animals. *All mice were housed in a 14 h (hours):10 h L(light):D(dark) cycle and we routinely used age-matched 12-to 16-week old male mice.* All mice had a C57BL/6J background. C57BL/6J, *Per2^-/-^* [Per2tm1Brd Tyrc-Brd/J (Zheng et al., 1999)], and PER2 luciferase [B6.129S6-*Per2^tm1Jt^*/J (Yoo et al., 2004)] mice were purchased from Jackson laboratories. *Per2^loxp/loxp^* and *Adorab2^loxp/loxp^* (Seo et al., 2015) mice were generated by Ozgene (Perth, Australia). VE-Cadherin-Cre [B6.Cg-Tg(Cdh5-cre)7Mlia/J(Alva et al., 2006)] or Germline Cre [B6.C-Tg(CMV-cre)1Cgn/J] were purchased from Jackson laboratories. To obtain endothelial tissue-specific mice, we crossbred *Per2^loxp/loxp^* mice with the VE-Cadherin-Cre recombinase mouse. To obtain *Adora2b^-/-^* mice we crossbred *Adorab2^loxp/loxp^* with the Germline Cre mouse. Before experiments, mice were housed for at least 4 weeks in a 14/10-h light-dark (lights on 6 AM [ZT0], lights off 8 PM[ZT14]) cycle to synchronize (entrain) the circadian clock of all mice to the ambient light-dark cycle. We conducted all mouse experiments at the same time points (ZT 3, ZT15), unless specified otherwise.

### Human subjects

Healthy human volunteers were exposed to intense light exposure (10,000 LUX) for 30 min every morning for five days from 8:30 AM – 9:00 AM. 5 mL blood was drawn on day one at 8:30 AM and 9:00 AM (before and after light exposure). We obtained approval from the Institutional Review Board (COMIRB #13-1607) for our human studies prior to written informed consent from everyone. A total of 17 healthy volunteers were enrolled (11 female and 6 male, age range between 21-44 yrs.).

## METHOD DETAILS

### Intense light exposure in mice

Mice were exposed to intense light (10,000 LUX, Lightbox, Uplift Technologies DL930, full spectrum, UV filter) for 3, 5 or 7 days and compared to mice maintained at room light [200 LUX] for 7 days (***Note****: infarct sizes are always the same at ZT 3 (9AM) regardless of the length of being housed under normal housing conditions*) (Bartman et al., 2017). To control for temperature changes, we measured the rectal temperature, using a rectal thermometer probe (RT, Effenberg, Munich, Germany).

### Cortisol measurements

To measure plasma cortisol levels in mice after one week of room light or intense light a mouse cortisol ELISA Kit (LifeSpan BioSciences, Inc., Seattle, WA) was used.

### Murine model for cardiac MI and heart enzyme measurement

Murine *in situ* myocardial ischemia and reperfusion injury (*60-min ischemia/120 min reperfusion)* and troponin-I (cTnI) measurements were performed as described (Eckle et al., 2006; Eckle et al., 2011; Eckle et al., 2007). Infarct sizes were determined by calculating the percentage of infarcted myocardium to the area at risk (AAR) using a double staining technique with Evan’s blue and triphenyltetrazolium chloride. AAR and the infarct size were determined via planimetry using the NIH software Image 1.0 (National Institutes of Health, Bethesda, MA). For troponin I (cTnI) measurements blood was collected by central venous puncture and cTnI was analyzed using a quantitative rapid cTnI assay (Life Diagnostics, Inc., West Chester, PA, USA). ***Note***: *cTnI is highly specific for myocardial ischemia and has a well-documented correlation with the infarct size in mice* (Eckle et al., 2006; Eckle et al., 2008; Eckle et al., 2007; Kohler et al., 2007) *and humans* (Vasile et al., 2008).

### Wheel running

Mice were maintained individually in running-wheel cages (Starr Life Sciences, wheel diameter: 11.5 cm). Running-wheel activity was recorded every 5 minutes using the Vital View Data Acquisition System (Starr Life Sciences, Oakmont, PA). Data were analyzed using BioDare (Biological Data Repository) (Moore et al., 2014). Amplitude of wheel running activity was calculated using the fast fourier transform non-linear least squares (FFT-NLLS) method. The distance walked was calculated as the sum of a 7 day-recording.

### Luciferase Assay

Expression of the PER2 protein using luciferase reporter mice was assayed as described (Eckle et al., 2012). Expression of the PER2 protein using luciferase reporter mice was determined using the tissue homogenates in T-per Tissue Protein Extraction Reagent (Pierce, Thermo Fisher Scientific, Waltham, MA). The homogenates were centrifuged for 30mins at 4900 x g at 4°C. The luciferase protein activity was measured by using the Dual-Luciferase Reporter Assay System from Promega according to the manufacturer’s instructions using a Biotek Synergy 2 Multimode Microplate Reader (Winooski, VT).

### Enucleation procedure in mice

Mice were pre-anesthetized with subcutaneous carprofen and buprenorphine injections and anesthetized with a ketamine/xylazine/ace cocktail. The periocular region was then clipped to remove surrounding hair. After a surgical scrub with betadine, the optic nerve and associated blood vessels were clamped with hemostats. After 2 minutes the entire globe of eye and the optic nerve were removed. For additional analgesia bupivacaine was dripped into the socket. Next, the complete upper and lower eyelid margin were removed with fine tip scissors and the eyelids were closed with 3-4 interrupted sutures. Mice were recovering in cages on warm water circulating blankets. Carprofen injections were repeated every 24 hours for 2 days post operatively. After a 2-week recovery period under standard housing conditions, mice underwent intense light housing or myocardial ischemia studies.

### ^13^C-tracers in vivo

C57BL/6 wildtype mice were housed under daylight (10,000 LUX, L:D 14:10 h) or standard room light (200 LUX, L:D 14:10 h) for 7 days followed by infusion of 10 mg/kg/min U-^13^C-glucose (Cambridge Isotope Laboratories, Tewksbury, MA) via an intra-arterial catheter over 90 minutes. Left ventricles of the hearts were flushed with ice cold KCl and the left ventricle was shock frozen and analyzed by liquid chromatography–tandem mass spectrometry. Isotope-resolved metabolite analyses were performed by LC-MS on a Waters Acquity ultrahigh-performance liquid chromatography (UPLC) system coupled to a Waters Synapt HDMS quadrupole time-of-flight mass spectrometer equipped with an atmospheric pressure electrospray ionization (ESI) source. LC-MS was performed with full-mass detection within m/z 100-1000 in the positive-ion or negative-ion mode as described below. The typical MS operation parameters included ESI spray voltage 2.8 kV, sampling cone voltage 4 V, drying gas flow 50 L/min / temperature 120 °C, and nebulizing nitrogen gas flow 700 L/h / temperature 350 °C. The authentic compound sodium D-fructose-6-sphosphate was obtained from Sigma-Aldrich and were used as the standard substances for location and identification of the targeted metabolites. UPLC conditions were optimized to ensure appropriate separation of each metabolite from its structural isomers and the interfering components detected in the samples. For sample preparation, frozen specimens were individually ground to fine powder in liquid nitrogen and were weighed to 5-mL borosilicate glass test tubes. Ice-cold methanol-water (50:50, v/v), equivalent to 1 mL per 100 mg tissue, was added to each tube. The samples were lysed on ice with 15 s x 2 sonications using a Fisher Scientific Model 100 cell dismembrator. After vortex-mixing and 5-min sonication in an ice-water bath, ice-cold methanol, equivalent to 1 mL per 100 mg tissue, was added. The sample tubes were capped, violently vortex-mixed for 2 min, sonicated in the same ice-water bath for 5 min and then placed at – 20 °C for 2 h. before the tubes were centrifuged at 4,000 rpm and 4 °C for 15 min in a Beckman Coulter Allegra X-22R centrifuge. The clear supernatant of each sample was collected and transferred to a 3-mL borosilicate glass test tube for the following LC-MS analyses. Fructose-6-phosphate was measured by ion-pairing LC-MS using tributylamine (TBA) as the paired counter ion reagent.(Luo et al., 2007) The chromatographic separation was conducted on a YMC-Triart C18 UPLC column (2.1 x 150 mm, 1.9 µm) using binary-solvent gradient elution with 2 mM TBA in water (pH adjusted to 6 with acetic acid) as mobile phase A and methanol as mobile phase B. The elution gradient was 0-9 min, 2% to 55 % B; 9-9.5 min, 55% to 100 % B; 9.5– 11 min, 100 % B. The column was equilibrated with 2% B for 5 min between injections. The column flow rate was 0.25 mL/min and the column temperature were 50 °C. A 100-µL aliquot of each supernatant from individual mouse heart specimens was dried under a nitrogen flow in a fume hood and the residue was reconstituted in 200 µL of mobile phase A. 5 µL was injected for LC-MS with negative-ion detection. The organic acids were analyzed by chemical derivatization LC-MS with 3-nitrophenyl hydrazine (3NPH) as the pre-analytical derivatizing reagent.(Han et al., 2013a) In brief, 100 μL of the supernatant was mixed 50 μL of 200 mM 3NPH.HCl in 75% methanol and 50 μL of 150 mM 1-ethyl-3-(3-dimethylaminopropyl)carbodiimide-HCl. The mixture could react at 30 °C for 30 min and was then mixed with 400 μL of water. 10-μL aliquots were injected onto a C8 UPLC column (2.1 x 50 mm, 1.7 μm) for LC-MS runs with negative-ion detection and using the LC procedure as described.(Han et al., 2013a) For all the above LC-MS analyses, the monoisotopic ion chromatograms of the metabolite, together with their isotopomeric counterparts resulting from the U-^13^C glucose tracer, were extracted based on their calculated m/z values, within a mass window of 60 ppm (+/-30 ppm), and their peak areas were integrated. The peak areas of any observed isotopomeric forms derived from the U-^13^C glucose tracer for the metabolite was corrected by subtracting the abundance contributions from the natural and any other U-^13^C glucose tracer-derived isotopic forms (Eckle et al., 2013; Eckle et al., 2012; Han et al., 2013a; Han et al., 2013b).

### PFK and cAMP-activity mouse tissue

To measure Phosphofructokinase activity from myocardium or plasma, C57BL/6J or *Per2^-/-^* mice were euthanized following light exposure. Blood was removed to obtain plasma further analysis, and the tissue was immediately flash frozen at -80°C. Enzyme activity from homogenized tissue or plasma was determined using a PFK Assay Kit from Biovison. To determine cAMP in cardiac tissue, a cyclic AMP Enzyme Immunoassay Kit (Mouse/Rat cAMP Parameter Assay Kit, R&D, Minneapolis, MN) was used according to the manufacturer’s protocol.

### Adenosine measurements

Whole murine hearts were collected in 1 mL of 80% MeOH and flash-frozen under liquid nitrogen and stored at -80°C. Adenosine was extracted and quantified in the tissue as described (Lee et al., 2018). Analyses were performed on an Agilent Technologies 1260 Infinity HPLC using a phenomenex Luna C18(2) column (100 Å, 150 X 4.6 mm) (mobile phase A: 50 mM KH2PO4, 5 mM tetrabutylammonium bisulfate, pH 6.25; mobile phase B: acetonitrile; column temperature: 30°C; flow rate: 1 mL/min; 75 µL injection). Samples were filtered through VIVASPIN 500 membranes (Sartorius Stedim Biotech, 5,000 MWCO, PES) prior to HPLC analysis. Chromatographic separation of the metabolites was performed using a combination of isocratic and gradient methods including column washing and equilibration periods at the end (0 min: 100% A; 7 min: 100% A; 10 min: 97% A; 18 min: 97% A; 45 min: 86% A; 60 min: 50% A; 80 min: 50% A; 90 min: 100% A; 135 min: 100% A). Adenosine was detected by absorption at 254 nm, and the absorbance spectra and retention time verified by co-injection with an authentic standard.

### HIF1A-HRE binding assay

Nuclear protein fractions were isolated from heart tissue using NE-PER kit following the manufacturer’s instructions (Thermo Fisher Scientific, Waltham, MA). Protein was quantified using the BCA Protein Assay Kit (Thermo Fisher Scientific, Waltham, MA). HIF1A transcription factor from 20 μg of nuclear protein was measured using a HIF1A Transcription Factor Assay kit (Abcam, ab133104, Cambridge, MA).

### Microarray analysis

Total RNA was isolated from heart tissue from intense light or room light ‘pretreated’ C57BL/6J or *Per2^-/-^* mice with the RNeasy micro kit (Qiagen, Valencia, CA) using Qiagen on-column DNase treatment to remove any contaminating genomic DNA. The integrity of RNA was assessed using a Tapestation 2200 (Agilent Technologies) and RNA concentration was determined using a NanoDrop ND-1000 spectrophotometer (NanoDrop, Rockland, DE). Hybridization cocktail was prepared starting with 100ng Total RNA using the GeneChip WT PLUS Reagent Kit. Samples were hybridized to the Arrays (Mouse Clariom D arrays) for 16 hours at 45 degrees C in a GeneChip Hybridization Oven 645. Arrays were then washed and stained in a GeneChip Fluidics Station 450 and scanned in a GeneChip Scanner 3000. Each array was subjected to visual inspection for gross abnormalities. Several other QC metrics were used to monitor hybridization efficiency and RNA integrity over the entire processing procedure. Raw image files were processed using Affymetrix GCOS 1.3 software to calculate individual probe cell intensity data and generate CEL data files. Using GCOS and the MAS 5.0 algorithm, intensity data was normalized per chip to a target intensity TGT value of 500 and expression data and present/absent calls for individual probe sets calculated. Gene symbols and names for data analyzed with the MAS 5.0 algorithm were from the Affymetrix NetAffx Mouse 430_2 annotations file. Quality control was performed by examining raw DAT image files for anomalies, confirming each GeneChip array had a background value less than 100, monitoring that the percentage present calls was appropriate for the cell type, and inspecting the poly (A) spike in controls, housekeeping genes, and hybridization controls to confirm labeling and hybridization consistency. According to our experimental setup the arrays were normalized, grouped and analyzed for differentially expressed transcripts based on different statistical tests. Different clustering algorithms allowed us to identify transcripts that show similar expression profiles. Using the TAC (Transcriptome Analysis Console, Affymetrix) software we were able to identify biological mechanisms, pathways and functions most relevant to our experimental dataset. Array data have been deposited in the Array Express database at EMBL-EBI (www.ebi.ac.uk/arrayexpress) under accession number E-MTAB-7196 (http://www.ebi.ac.uk/arrayexpress/experiments/E-MTAB-7196).

### Modified Miles Assay

Mice were injected with Evans blue dye (80mg/kg) via a carotid catheter at the beginning of reperfusion after 60 minutes of ischemia. After 2 hours of reperfusion, mice were euthanized and perfused with citrate buffer, pH4. The hearts were then excised, cut into 1mm thick slices and photographed. Heart slices were then incubated in 1ml of formamide overnight at 70 °C. After centrifugation, absorbance at 620 nm was measured by using a spectrophotometer Biotek [Synergy 2 Multimode Microplate Reader (Winooski, VT)]. Extravasated Evans blue (ng) was determined from a standard curve and normalized to tissue weight (g). Evans blue dye content was determined from standard curve and normalized to heart tissue weight (Galaup et al., 2012).

### Chromatin immunoprecipitation (ChIP) assay from mouse heart tissue

ChIP assays for heart tissue were performed using the SimpleChIP® Plus Enzymatic Chromatin IP Kit (Cell signaling). Briefly, freshly whole hearts were placed into PBS containing 1.5% formaldehyde and homogenized using a probe tissue homogenizer. Crosslinking was stopped after 15 min by adding glycine for 5min. The samples were centrifuged to collect the pellet. Pellets were disaggregated with a dounce homogenizer and chromatin was enzymatically digested to yield 200- to 1,500-bp DNA fragments. The chromatin was incubated at 4°C overnight with a HIF1A antibody (NB100-134, Novus, Littleton CO) or rabbit IgG control (Cell Signaling, Danvers, MA). After reverse cross-linking by heating the samples at 65°C overnight and treating with Proteinase K, DNA was purified using DNA purification kit (Cell Signaling, Danvers, MA). Quantitative analyses of DNA products obtained from ChIP assay were performed by real-time RT-PCR with primers specific for the mouse ANGPTL4 promoter. The ChIP data were normalized against IgG to account for non-specific immunoprecipitation (fold enrichment relative to the negative (IgG) sample). Primers used were: Primer 1: sense CCC CAC TTG CCA TCT GAA CT, antisense GAT GCC TTC TTG ACT GAC CCC, Primer 2: sense GGG AAT TTC CGG CCT TAG GAT, antisense GTT CTT GGG GAT GGC TGC TTC, Primer 3: sense CCT GGG ACG AGA TGA ACT TGC, antisense ATC TTT TCC CTT GGG CCC CT for the mouse ANGPTL4 promoter (see also Figure S3).

### Hypoxia exposure

For hypoxia experiments cells were placed in a hypoxia chamber (Coy Laboratory Products Inc., Grass Lake, MI) in preequilibrated hypoxic medium at 1% O2 for 24 hours.

### Lentiviral-mediated generation of cells with knockdown of PER2 or HIF1A

Stable cell cultures with decreased PER2 and HIF1A expression were generated by lentiviral-mediated shRNA expression. pLKO.1 lentiviral vector targeting PER2 had shRNA sequence of CCG GGA CAC ACA CAA AGA ACT GAT ACT CGA GTA TCA GTT CTT TGT GTG TGT CTT TTT (TRCN0000018542) and HIF1A had shRNA sequence of CCG GCC AGT TAT GAT TGT GAA GTT ACT CGA GTA ACT TCA CAA TCA TAA CTG GTT TTT (TRCN 0000003809). For controls, nontargeting control shRNA (SHC002; Sigma) was used. HMEC-1 were co-transfected with pLK0.1 vectors and packaging plasmids to produce lentivirus. Filtered supernatants were used for infection of HMEC-1 and cells were selected with puromycin or geneticin until a knockdown was confirmed (Eckle et al., 2013).

### Transcriptional Analysis

Total RNA was isolated using Trizol Reagent (Invitrogen, Carlsbad, CA), phenol-chloroform extraction, and ethanol precipitation followed by purification in a modified protocol from RNeasy Mini Kit (Qiagen, Germantown, MD). RNA was quantified using either a Qubit 3.0 RNA BR Assay Kit (Thermo Fisher Scientific, Waltham, MA) or Nanodrop 2000. Quantification of transcript levels was determined by real-time RT-PCR (iCycler; Bio-Rad Laboratories, Inc, Hercules, CA). qPCR reactions contained 1x final primer concentration (Qiagen primers, Germantown, MD) or 1 µM sense and 1 µM antisense oligonucleotides (Invitrogen custom DNA oligos, Carlsbad, CA) with SYBR Green (Bio-Rad, Hercules, CA). Primer sets used (Qiagen QuantiTect) were Hs_PER2_1_SG (QT0011207), Hs_PKM_1_SG (QT00028875), Hs_LDHA_1_SG (QT00001687), Hs_SIRT3_1_SG (QT00091490), Hs_ACTB_2_SG (QT01680476), and Hs_COX4|2_1_SG (QT00044933), Mm_Per2_1_SG (QT00198366), Mm_Angptl4_1_SG (QT00139748), Mm_Cox4i2_1_SG (QT00137844), Mm_Cldn1_1_SG (QT00159278), Mm_Actb_2_SG (QT01136772). Primers for human OPN4 (Invitrogen, Carlsbad, CA, sense 5’-AGT CGC CCC TAC CCC AGC TA-3’ and antisense 5’-CAC AGC TGC TGC CTC CAT GT-3’) were custom designed. Each target sequence was amplified using the following protocol: 95°C for 3 min, 40 cycles of 95°C for 15 sec, 55°C for 0.5 min, 72°C for 10 sec, 72°C for 1 min and a melt curve protocol.

### Immunoblotting experiments

Protein was isolated from HMEC-1 using M-Per following manufacturer’s instructions (Thermo Fisher Scientific, Waltham, MA) and including a protease and phosphatase inhibitor cocktail (Thermo Fisher Scientific, Waltham, MA). Protein was quantified using a Qubit Fluorometer 3.0 and Qubit Protein Assay Kit (Thermo Fisher Scientific, Waltham, MA). 5 – 25 μg of protein was denatured at 95°C in Laemmli sample buffer for 5 min. Samples were resolved on a 4 – 10% polyacrylamide gel and transferred to nitrocellulose membranes, which were blocked for 1h at room temperature in either 5% BSA / TBST or 5% milk / TBST. The membranes were incubated in primary antibody at a concentration of 1:1000 overnight at 4°C. The primary antibodies used were rabbit polyclonal PER2 (Novus Biologicals, NB100-125, Littleton CO, or Abcam, ab64460, Cambridge, MA), mouse monoclonal actin (Ab-1) (JLA20, Calbiochem, Diego, CA,), rabbit polyclonal IDH2 (Novus Biologicals, NBP2-22166, Littleton CO), rabbit polyclonal SUCLG1 (Novus Biologicals, NBP1089489, Littleton CO), rabbit polyclonal ACO2 (Novus Biologicals, H00000050-D01P, Littleton CO), rabbit polyclonal SIRT3 (Abcam, ab86671, Cambridge, MA), anti-alpha Tubulin antibody (Abcam, ab7291, Cambridge, MA), Anti-VDAC1 / Porin antibody (Abcam, ab15895, Cambridge, MA), Anti-TATA binding protein (TBP) antibody (Abcam, ab51841, Cambridge, MA), mouse monoclonal β-ACTIN (Cell Signaling Technologies, 8H10D10, Danvers, MA), and mouse monoclonal anti-DDK (FLAG) (OriGene Technologies, TA50011-100, Rockville, MD). The next day, blots were washed 3 – 4x with TBST and incubated with secondary antibody at a concentration of 1:5000 in the respective blocking buffer, washed an additional 3 times, and visualized using SuperSignal West Femto Maximum Sensitivity Substrate (Thermo Fisher Scientific, Waltham, MA). The secondary antibodies used were goat polyclonal IgG (Novus Biologicals, NB7487), goat anti-mouse IgM (Calbiochem, San Diego, CA), and goat anti-rabbit IgG (Thermo Fisher Scientific, Waltham, MA).

### Lactate measurements

Lactate measurements were done using the L-Lactate Colorimetric Assay Kit following the manufacturer’s protocol (Abcam, Cambridge, MA).

### Cytotoxicity

Cytotoxicity was determined using the LDH-Cytotoxicity Assay Kit per manufacturer’s protocol (Abcam, Cambridge, MA).

### Seahorse stress tests

*Glycolytic stress tests.* The XF24 Seahorse Bioanalyzer was used in conjunction with glycolytic stress tests following manufacturer’s specifications (Agilent, Santa Clara, CA). Cells were plated in the morning at a density of 1.2×10^5^ cells / well and serum starved in the evening one day prior to assaying. One hour prior to the stress test cells were incubated in XF Assay Medium (Agilent, Santa Clara, CA) at a pH of 7.4. Final concentration of glucose was 10 mM / well, oligomycin 1.0 μM / well, and 2-deoxyglucose 50 mM / well. *Mitochondrial stress tests.* The XF24 Seahorse Bioanalyzer was used for mitochondrial stress tests (Agilent, Santa Clara, CA). For all assays, pH and oxygen consumption rate (OCR) were measured. For TCA cycle readouts, an additional measurement of carbon dioxide evolution rate (CDER) was measured. Final concentrations were 1.0 μM oligomycin, 3.6 μM FCCP, and 1.125 μM Rotenone / Antimycin A. *Cell energy phenotype assays.* Following manufacturer’s instructions (Agilent, Santa Clara, CA), the Seahorse Bioanalyzer was used to assess cell energy phenotype at baseline.

### Chromatin immunoprecipitation (ChIP) assay-cell culture

ChIP assays were performed using the ChIP-IT™ Express Enzymatic Kit from Active Motif (Carlsbad, CA, USA). Briefly, Scr and PER2KD HMECs were grown to 90% confluence in phenol red-free Dulbecco’s modified Eagle medium (DMEM) supplemented with 10% charcoal DEXTRAN-stripped FBS for at least 3 days. After hypoxia exposure at 1% O2 for 24h, ChIP assays were performed according to manufacturer’s protocol. Briefly, chromatin was cross-linked in 1% formaldehyde in minimal cell culture medium (Invitrogen, Carlsbad, CA), and nuclei were extracted. Chromatin was enzymatically digested for 11mins to yield 200- to 1,500- bp DNA fragments and the supernatant containing precleared chromatin was then incubated at 4°C overnight with mouse monoclonal HIF1A antibody (H1alpha67, ChIP Grade, Abcam, Cambridge, MA) or rabbit IgG control (Cell Signaling, Danvers, MA). After reverse cross-linking by heating the samples at 65°C overnight and treating with Proteinase K, DNA was purified using phenol-chloroform extraction. Quantitative analyses of DNA products obtained from ChIP assay were performed by RT-PCR with primers specific for the human LDHA promoter. RT-PCRs conducted on DNA derived from input chromatin templates served as positive controls whereas reactions conducted on IgG-precipitated templates served as negative controls. The RT-PCR signal was barely detectable for these controls. The signal for these samples and IgG-precipitated templates was negligible on gels. Primers used were: sense ATT ACG TGC CAG AAG CTG TT and antisense TTT CCT CAT CCA TGA AAC CT for the human LDHa promoter. Conventional PCR signals were stained with ethidium bromide in 1% agarose gels.

### Affinity purification-mass spectrometry-based proteomics

HMEC-1 were placed in a hypoxia chamber (Coy Laboratory Products Inc., Grass Lake, MI) in preequilibrated hypoxic medium at 1% O2. Following 24 h of hypoxia, the samples were isolated for cytoplasmic and nuclear protein fractions according to the NE-PER kit specifications (Thermo Fisher Scientific, Waltham, MA). To identify interacting proteins with PER2, co-immunoprecipitation (Co-IP) for PER2 was performed using the Pierce Co-IP Kit (Thermo Fisher Scientific, Waltham, MA). Specifically, 10 μg of rabbit anti-PER2 antibody (Novus, NB100-125) was immobilized to the amine-reactive resin. 100 μg of sample was incubated overnight at 4 °C with the anti-PER2 coupled resin. Samples were washed and then eluted. Samples were then loaded onto a 1.5 mm thick NuPAGE Bis-Tris 4-12% gradient gel (Invitrogen, Carlsbad, CA). The BenchMark™ Protein Ladder (Invitrogen, Carlsbad, CA) was used as a protein molecular mass marker. The electrophoretic run was performing by using MES SDS running buffer, in an X-Cell II mini gel system (Invitrogen) at 200 V, 120 mA, 25 W per gel for 30 minutes. The gel was stained using SimplyBlue™ SafeStain (Invitrogen, Carlsbad, CA) stain and de-stained with water according to the manufacturer’s protocol. Each lane of the gel was divided into 9 equal-sized bands, and proteins in the gel were digested as follows. Gel pieces were destained in 200 µL of 25 mM ammonium bicarbonate in 50 % v/v acetonitrile for 15 min and washed with 200 µL of 50% (v/v) acetonitrile. Disulfide bonds in proteins were reduced by incubation in 10 mM dithiothreitol (DTT) at 60 °C for 30 min and cysteine residues were alkylated with 20 mM iodoacetamide (IAA) in the dark at room temperature for 45 min. Gel pieces were subsequently washed with 100 µL of distilled water followed by addition of 100 mL of acetonitrile and dried on SpeedVac (Savant, Thermo Fisher Scientific, Waltham, MA). Then 100 ng of trypsin was added to each sample and allowed to rehydrate the gel plugs at 4 °C for 45 min and then incubated at 37 °C overnight. The tryptic mixtures were acidified with formic acid up to a final concentration of 1%. Peptides were extracted two times from the gel plugs using 1% formic acid in 50% acetonitrile. The collected extractions were pooled with the initial digestion supernatant and dried on SpeedVac (Savant, Thermo Fisher Scientific, Waltham, MA). Samples were desalted on Thermo Scientific Pierce C18 Tip. For mass spectrometry, samples were analyzed on an LTQ Orbitrap Velos Pro mass spectrometer (Thermo Fisher Scientific, Waltham, MA) coupled to an Eksigent nanoLC-2D system through a nanoelectrospray LC − MS interface. A volume of 8 μL of sample was injected into a 10 μL loop using the autosampler. To desalt the sample, material was flushed out of the loop and loaded onto a trapping column (ZORBAX 300SB-C18, dimensions 5×0.3 mm 5 μm) and washed with 0.1% FA at a flow rate of 5 μL/min for 5 min. The analytical column was then switched on-line at 600 nl/min over an in house-made 100 μm i.d. × 150 mm fused silica capillary packed with 4 μm 80 Å Synergi Hydro C18 resin (Phenomex; Torrance, CA). After 10 min of sample loading, the flow rate was adjusted to 350 nL/min, and each sample was run on a 90-min linear gradient of 2–40% ACN with 0.1% formic acid to separate the peptides. LC mobile phase solvents and sample dilutions used 0.1% formic acid in water (Buffer A) and 0.1% formic acid in acetonitrile (Buffer B) (Optima™ LC/MS, Fisher Scientific, Pittsburgh, PA). Data acquisition was performed using the instrument supplied Xcalibur™ (version 2.1) software. The mass spectrometer was operated in the positive ion mode. Data acquisition was performed using the instrument supplied Xcalibur™ (version 2.1) software. The mass spectrometer was operated in the positive ion mode. Full MS scans were acquired in the Orbitrap mass analyzer over the *m/z* 350–1800 range with resolution 60,000 (*m/z* 400). The target value was 5.00E+05. The twenty most intense were selected for sequencing and fragmented in the ion trap with normalized collision energy of 35%, activation q = 0.25, activation time of 20 ms, and one microscan. The target value was 1.00E+04. The maximum allowed ion accumulation times were 500 ms for full scans and 150 ms for CID. For database searching and protein identification, MS/MS spectra were extracted from raw data files and converted into mgf files using MassMatrix (Cleveland, OH). These mgf files were then independently searched against mouse SwissProt database using an in-house Mascot™ server (Version 2.2, Matrix Science). Mass tolerances were +/-15 ppm for MS peaks, and +/-0.6 Da for MS/MS fragment ions. Trypsin specificity was used allowing for 1 missed cleavage. Met oxidation, protein N-terminal acetylation, peptide N-terminal pyroglutamic acid formation were allowed for variable modifications while carbamidomethyl of Cys was set as a fixed modification. Scaffold (version 4.4.6, Proteome Software, Portland, OR, USA) was used to validate MS/MS based peptide and protein identifications. Peptide identifications were accepted if they could be established at greater than 95.0% probability as specified by the Peptide Prophet algorithm. Protein identifications were accepted if they could be established at greater than 99.0% probability and contained at least two identified unique peptides. Following identification of potential PER2 interacting proteins in normoxia, hypoxia, and normoxia vs. hypoxia, lists obtained from Scaffold were analyzed by Ingenuity Pathway Analysis (Qiagen), Panther Classification System, and Reactome Analysis to detect pathways PER2 regulates in normoxia and hypoxia.

### Co-immunoprecipitations (Co-IPs)

Co-IPs were done using the Pierce Co-Immunoprecipitation (Co-IP) Kit (Thermo Fisher Scientific, Waltham, MA). 10 μg of antibody was immobilized to columns. Pulled-down protein was quantified using a Qubit Fluorometer 3.0 and resolved by immunoblotting as described.

### Subcellular compartment analysis

HMEC-1 were placed in hypoxia (1% O2) or normoxia for 24 h. After normoxia or hypoxia exposure, samples were isolated for cytoplasmic and nuclear protein fractions according to the NE-PER kit specifications (Thermo Fisher Scientific, Waltham, MA). Mitochondria protein fraction was isolated with the Dounce homogenization method according to the Mitochondria Isolation Kit for Cultured Cells specifications (Thermo Fisher Scientific, Waltham, MA).

### Immunocytochemistry and analysis of PER2 localization to mitochondria

Scr and PER2KD HMEC-1 cells were plated onto collagen coated cover slips and allowed to reach confluency. Following 24hrs of normoxia or hypoxia mitochondria were labeled using MitoTracker-deep red (100nM; Invitrogen) diluted in serum free media for 30mins at 37°C. Cells were then fixed with 4% PFA for 15mins at 37°C. Following PBS washes, samples were incubated with rabbit anti-Per2 (1:5000; Millipore) with 2% lamb serum and 0.1% Triton-X overnight at 4°C. Cells were then washed with PBS and incubated with Alexa Fluor goat-anti rabbit IgG-488 (1:500; Invitrogen) and DAPI (1:2000; Invitrogen). Following PBS washes, the cover slips were mounted for imaging. Confocal immunofluorescent images were captured using a Zeiss 780 LSM confocal microscope with gain and laser power remaining constant between all samples. Samples were randomized and three images were blindly taken from each sample (n=6/condition). Zen imaging software was used to assess Per2 fluorescent intensity and normalized to the total number of cells per 40x field. Colocalization of Per2 to mitochondria was assessed using ImageJ colocalization threshold analysis with Pearson’s correlation coefficient being reported. Analysis of the three images from each sample was averaged for each sample. Representative images were chosen for figures to best depict the results obtained.

### Enzyme activities IDH, ACO, SUCLG, Complex IV, PFK

Human isocitrate dehydrogenase (IDH, Biovision, Milpitas, CA), aconitase (ACO, Abcam, Cambridge, MA), succinyl-CoA synthetase (SUCLG, Abcam, Cambridge, MA), phosphofructokinase (PFK, Biovision, Milpitas, CA) or mouse complex 4 activity (Abcam, Cambridge, MA) were measure colorimetric assay kits adhering to manufacturer’s instructions.

### Mitochondrial membrane potential dyes

MitoTracker Red CMXRos (Invitrogen Molecular Probes, Carlsbad, CA) and JC-1 Mitochondrial Membrane Potential Assay (Abcam, Cambridge, MA) were used per manufacturer’s specifications using 5 μM of JC-1 for 30 minutes at 37C. JC-1 quantification was done by calculating mean intensity.

### ^13^C-tracers in vitro

HMEC-1s were serum starved in either MCDB131 (low glucose) or glucose-free DMEM for 24 h prior to assay. Respective mediums were supplemented with either 12 mM U-^13^C-glucose (Cambridge Isotope Laboratories, Tewksbury, MA) or 166.67 μM 1,2-^13^C2-palmitic acid (Cambridge Isotope Laboratories, Tewksbury, MA) in hypoxia or normoxia for 24 h. Frozen cell pellets were extracted at 2e^6^ cells/mL in ice cold lysis/ extraction buffer (methanol:acetonitrile:water 5:3:2). Samples were agitated at 4 °C for 30 min followed by centrifugation at 10,000 g for 10 min at 4 °C. Protein and lipid pellets were discarded, and supernatants were stored at -80 °C prior to metabolomics analysis. Ten µL of extracts were injected into a UHPLC system (Vanquish, Thermo, San Jose, CA, USA) and run on a Kinetex C18 column (150 x 2.1 mm, 1.7 µm - Phenomenex, Torrance, CA, USA). Solvents were Optima H2O (Phase A) and Optima acetonitrile (Phase B) supplemented with 0.1% formic acid for positive mode runs and 1 mM NH4OAc for negative mode runs. For [U-^13^C]-glucose flux analysis (Thwe et al., 2017), samples were run on a 3 min isocratic (95% A, 5% B) run at 250 µL/min (Nemkov et al., 2015; Nemkov et al., 2017). For [1,2-^13^C2]-palmitate flux analysis, samples were analyzed using a 9 min gradient from 5-95% acetonitrile organic phase at 400 µL/min. The autosampler was held at 7 °C for all runs; the column compartment was held at 25 °C for the 3 min method and 45 °C for the 9 min method (McCurdy et al., 2016). The UHPLC system was coupled online with a Q Exactive mass spectrometer (Thermo, Bremen, Germany), scanning in Full MS mode (2 µscans) at 70,000 resolution in the 60-900 m/z range in negative and then positive ion mode (separate runs). Eluate was subjected to electrospray ionization (ESI) with 4 kV spray voltage. Nitrogen gas settings were 15 sheath gas and 5 auxiliary gas for the 3 min runs; 45 sheath gas and 15 auxiliary gas for the 9 min runs. Metabolite assignments and isotopologue distributions were determined using Maven (Princeton, NJ, USA), upon conversion of ‘*.raw’* files to ‘.*mzXML*’ format through MassMatrix (Cleveland, OH, USA). Chromatographic and MS technical stability were assessed by determining CVs for heavy and light isotopologues in a technical mixture of extract run every 10 injections.

### Cell permeability - TEER method

HMEC-1 were grown on polycarbonate permeable supports (0.4-μm pore, 6.5-mm diam; Corning Life Sciences, Acton, MA) at a cell density of 0.3 × 10^4^ cells / well. Confluent HMEC-1 were placed in a hypoxia chamber (Coy Laboratory Products Inc., Grass Lake, MI) in preequilibrated hypoxic medium at 1% O2. Transendothelial electrical resistance (TEER) was measured with the EVOM2 voltohmmeter (World Precision Instrument, Sarasota, FL, USA). Value of blank well was subtracted.

### Light sensing cells

HMEC-1 WT cells were transfected with pCMV6-Entry (C-terminal Myc and DDK Tagged, OriGene Technologies, Rockville, MD) or OPN4 (Myc-DDK-Tagged)-pCMV6-Entry transcript variant 1 (TrueORF Gold Expression-Validated cDNA Clones from OriGene Technologies, Rockville, MD) using FuGene HD Transfection Reagent (Promega, Madison, WI). After transfection, cells were kept in complete darkness until room light (∼200 LUX) or intense light (∼10,000 LUX) exposures for 30 minutes. Melanopsin protein expression validation was done by isolating protein using RIPA buffer (Thermo Fisher Scientific, Waltham, MA) supplemented with protease and phosphatase inhibitor cocktail (Thermo Fisher Scientific, Waltham, MA). Immunoblotting for anti-DDK (OriGene Technologies, Rockville, MD) was used to detect DDK-tagged melanopsin in transfected cells. Light-sensing cells subjected to glycolytic or mitochondrial stress tests on the Seahorse Bioanalyzer were exposed to light 30 min prior to Seahorse analyses.

### cAMP ELISA and phospho-CREB assays

Phospho-CREB (S133) Immunoassay (R&D Systems, Minneapolis, MN) or cAMP Parameter Assay Kit for human (R&D Systems, Minneapolis, MN) were used according to the manufacturer’s protocol.

### Human light exposure

Healthy human volunteers were exposed to intense light exposure (10,000 LUX) for 30 min every morning for five days from 8:30 AM – 9:00 AM. 5 mL blood was drawn on day one at 8:30 AM and 9:00 AM (before and after light exposure). While light exposure was repeated every morning for the five days, the next blood draws were on day two, three and five at 9:00 AM as indicated. In subset of experiments blood draws were also performed at 9 PM after 5 days of intense light therapy. For experiments involving actigraphy watches, the same volunteers wore the watch for one week without intense light exposure and maintained wearing the watches during the second week when light exposure was included. Actigraphy data were obtained by using a validated accelerometer (Actiwatch 2). We obtained approval from the Institutional Review Board (COMIRB #13-1607) for our human studies prior to written informed consent from everyone. A total of 17 healthy volunteers were enrolled (11 female and 6 male, age range between 21-44 yrs.).

### Human plasma melatonin, HIF1A and triglyceride levels

Melatonin levels were measured using the MT Elisa Kit for humans (My BioSource, San Diego, CA). HIF1A levels from human plasma samples were measured using the human HIF1A ELISA Kit from Invitrogen (Carlsbad, CA). Triglycerides were determined using a human Triglyceride Quantification Assay kit (Abcam, Cambridge, MA).

### Targeted metabolomics - mass spectrometry

Targeted metabolomics of human plasma following light exposure was performed as previously reported (A three-minute method for high-throughput quantitative metabolomics and quantitative tracing experiments of central carbon and nitrogen pathways (Nemkov et al., 2017). In brief, plasma samples were diluted at 1:25 in ice cold extraction solution (methanol, acetonitrile, water at a ratio of 5:3:2) and vortexed for 30 minutes at 4° C followed by removal of insoluble proteins and lipids by centrifugation at 10,000 RCF for 10 minutes at 4° C. The supernatants were collected and stored at -80° C until analysis. Analyses were performed using a Vanquish UHPLC system coupled to a Q Exactive mass spectrometer (Thermo Fisher Scientific, San Jose, CA, USA). Samples were resolved by a Kinetex C18 column (2.1 × 150 mm, 1.7 μm particle size; Phenomenex, Torrance, CA, USA) with a guard column at 25°C with an isocratic condition of 5% acetonitrile, 95% water, and 0.1% formic acid with a flow rate of 250 μL/min. The mass spectrometer was operated independently in positive or negative ion mode, scanning in Full MS mode from 60 to 900 m/z at 70,000 resolutions, with 4 kV spray voltage, 15 sheath gas, 5 auxiliary gas. Calibration was performed prior to analysis. Acquired data was converted from. raw to. mzXML file format using Mass Matrix (Cleveland, OH, USA). Metabolite assignments, isotopologue distributions, and correction for expected natural abundances of deuterium, 13C, and 15N isotopes were performed using MAVEN (Princeton, NJ, USA). Metabolite assignments were referenced to experimental retention times for over 400 analytical standards (MSMLS, IROATech, Bolton, MA, USA) and were determined over a 3-minute isocratic method with 20 μL of standards and samples injected for UHPLC/MS analysis.

## QUANTIFICATION AND STATISTICAL ANALYSIS

For comparison of two groups the unpaired student t-test was performed. For multiple group comparisons a one-way analysis of variance with a Tukey’s post hoc test was performed. Values are expressed as mean±SD. Assay details and n numbers are specified in the figure legends. P<0.05 was considered statistically significant. Only two-sided tests were used, and all data analyzed met the assumption for the specific statistical test that was performed. For all statistical analysis, GraphPad Prism 7.0 software was used. The authors had full access to and take full responsibility for the integrity of the data. All authors have read and agree to the manuscript as written.

## DATA AND CODE AVAILABILITY

The Array data have been deposited in the Array Express database at EMBL-EBI (www.ebi.ac.uk/arrayexpress) under accession number E-MTAB-7196 (http://www.ebi.ac.uk/arrayexpress/experiments/E-MTAB-7196

---

**Figure S1. Related to Figure 1.**
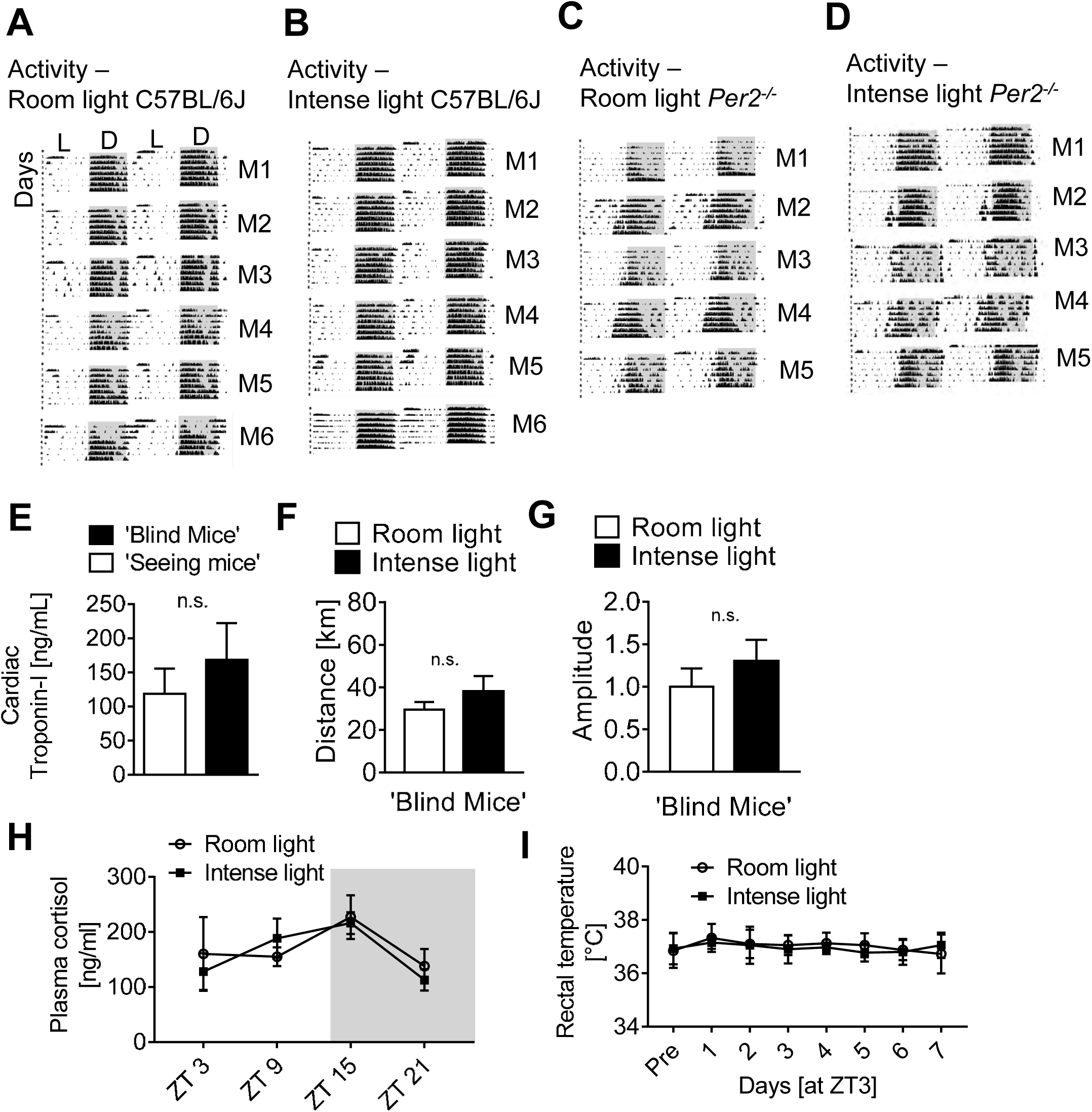
(**A-D**) 7 day wheel running activity graphs (double-plotted) from wildtype or *Per2^-/-^* mice exposed to 7 days room light versus 7 days intense light (mean±SD, n=5-6, M=mouse, L=light phase, D=dark phase, double-plotted actograms, ***Note***: numbers on the left indicate days. (**E**) Cumulative cardiac troponin measurements (ZT3+ZT15) from ‘seeing’ compared to enucleated ‘blind’ wildtype mice subjected to 60 min ischemia and 2 h reperfusion (mean±SD, n=7); (**F-G**) Wheel running measurements during 7 days of room light or intense light housing conditions in ‘blind’ C57BL/6J mice (mean±SD; n=4). **(H**) Plasma cortisol levels after 7 days of room or intense light exposure in C57BL/6J mice (mean±SD, n=5, (**I**) Rectal temperatures during 7 days of room light or intense light at ZT3 in C57BL/6J mice (mean±SD; n=4).

**Figure S2. Related to Figure 3.**
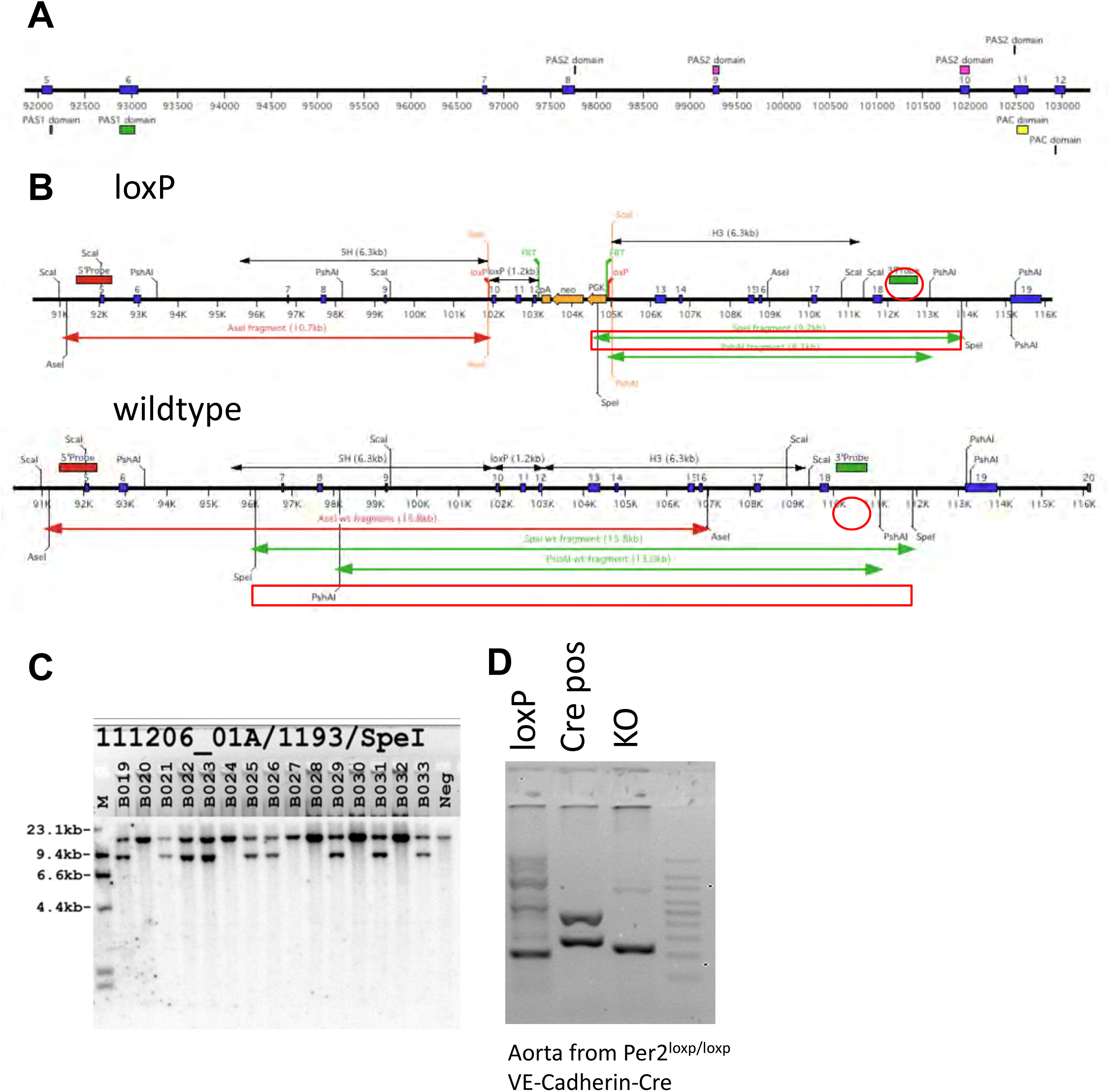
(**A**) *Per2^loxp/loxp^* -strategy: deletion of exons 10, 11 and 12 in the *Per2* gene removes half of the PAS2 domain and all of the PAC domain. This deletion also results in a frameshift mutation introducing an early stop codon. (**B**) Screening strategy. (**C**) Screening: DNA was digested with SpeI and probed with the P3 probe. These mice were the result of a wt (wildtype) /loxP x C57BL/6J mating. The expected sizes were: wildtype 15.8kb and loxP 9.2kb. Correct integration was confirmed by full sequencing. (**D**) PCR-Genotyping of aortic tissue from a *Per2^loxP/loxP^* -VE-Cadherin-Cre mouse.

**Figure S3. Related to Figure 3.**
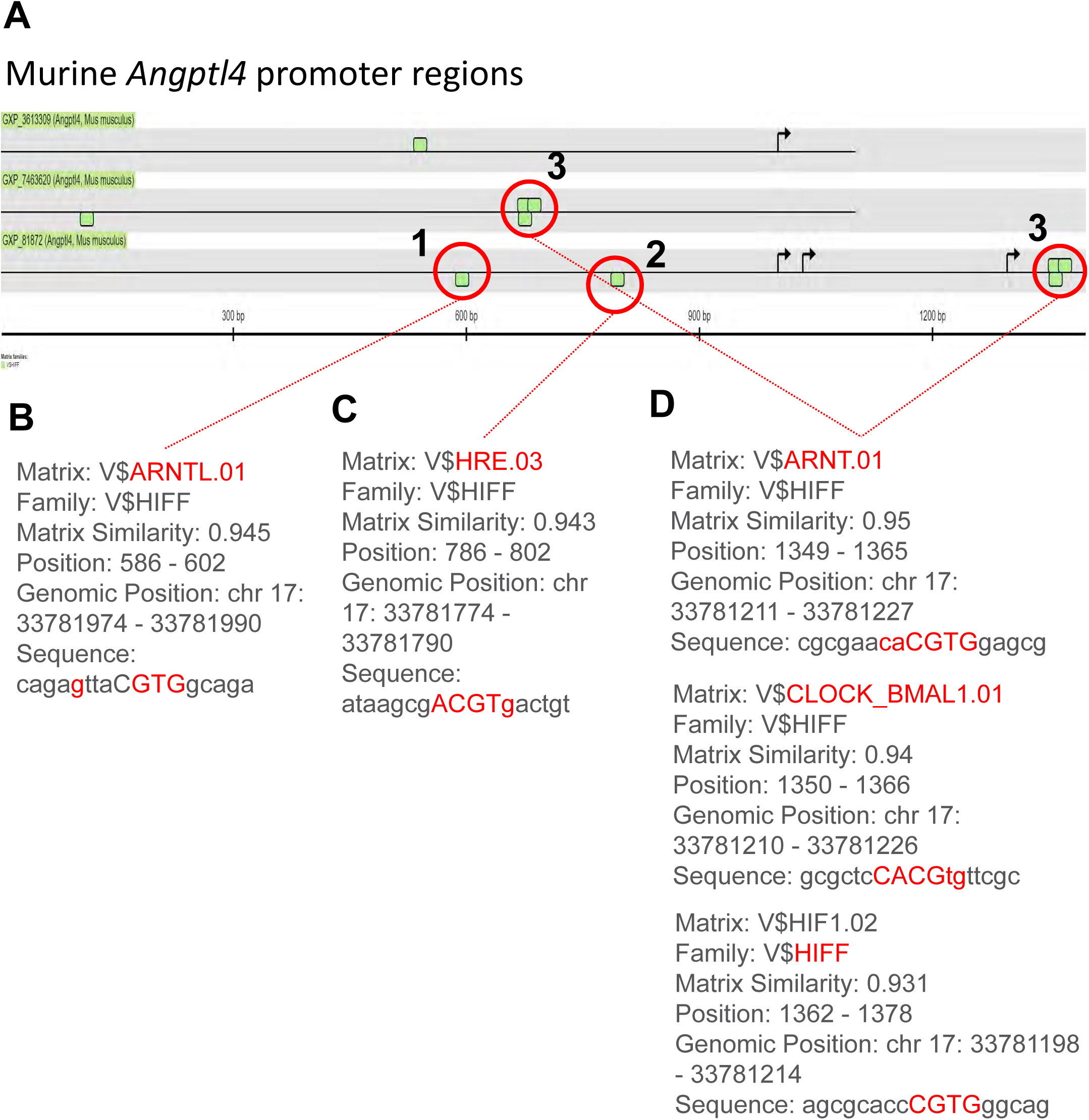
(**A**) Mouse *Anptl4* promotor regions (Genomatix). HIFF family binding sites are depicted by green boxes. (**B-D**) Primers for the ChIP assay covered the regions marked with a red circle (*Angptl4*-HRE1-3, **see Figure 3)**.

**Figure S4. Related to Figure 4.**
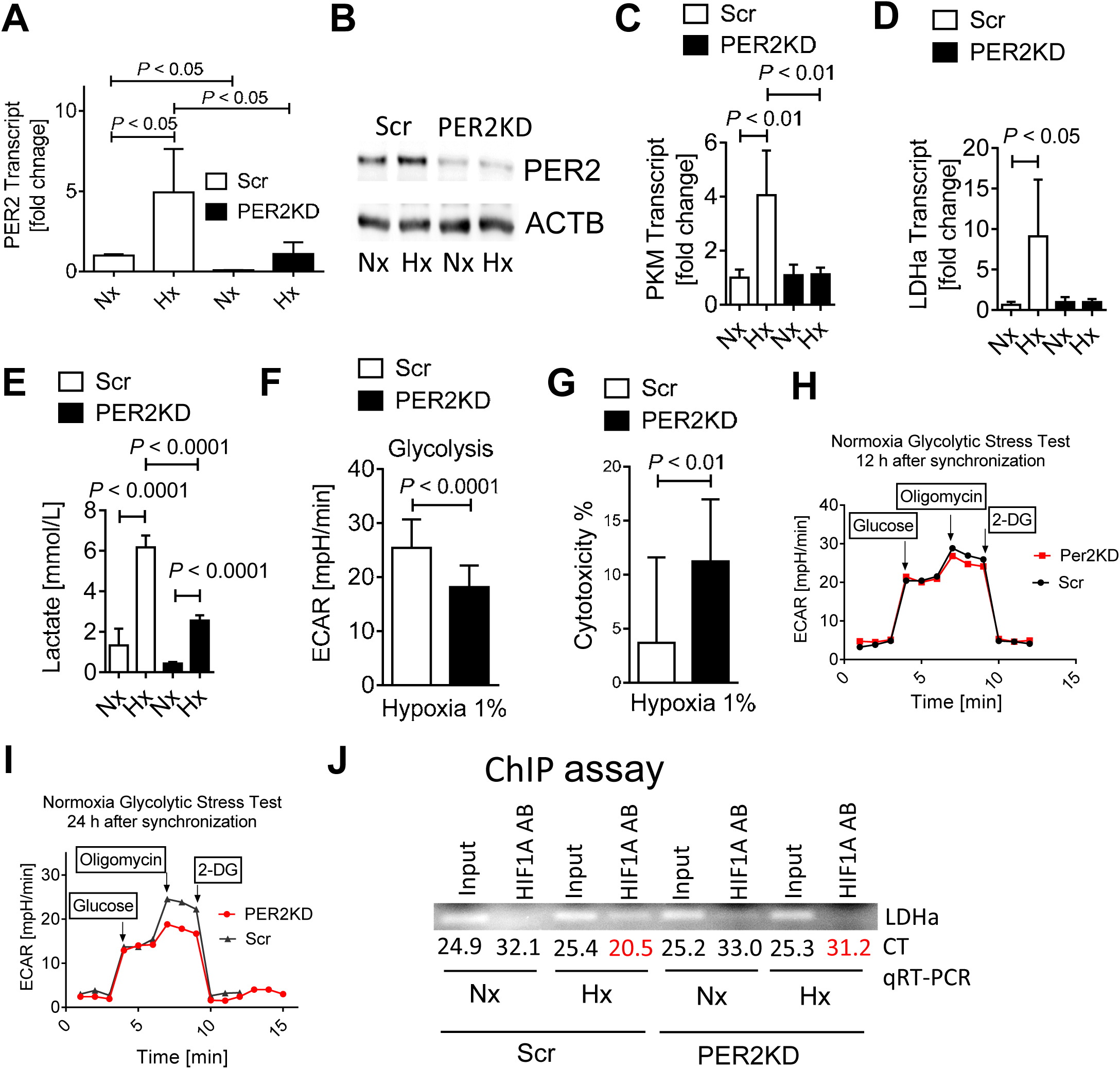
HMEC-1 controls (Scr; treated with lentiviral scrambled shRNA) or HMEC-1 PER2 knockdown (KD; treated with lentiviral PER2 shRNA) were synchronized via serum starvation and exposed to 24 h of normoxia (Nx) or 1% hypoxia (Hx) in all experiments unless specified otherwise. (**A, B**) PER2 transcript or protein levels (mean±SD, n=3). (**C, D**) Transcript expression of pyruvate kinase (PKM) or lactate dehydrogenase (LDHa) (mean±SD, n=3). (**E**) Lactate levels in cell supernatants (mean±SD, n=3). (**F**) Glycolytic stress test (mean±SD, n=10). (**G**) LDH-Cytotoxicity (mean±SD, n=10). (**H,I**) Glycolytic stress tests 12 or 24h after cell synchronization (mean±SD, n=10) under normoxic conditions. (**J**) Chromatin immunoprecipitation (ChIP) analysis to detect HIF1A protein binding to the human LDHa promoter. qRT-PCR for the human LDHa promoter was performed for quantification. PCR products analyzed on a 2% agarose gel (top, **not** quantitative) or quantitative CT values form the qRT-PCR are shown (bottom, n=3).

**Figure S5. Related to Figure 4.**
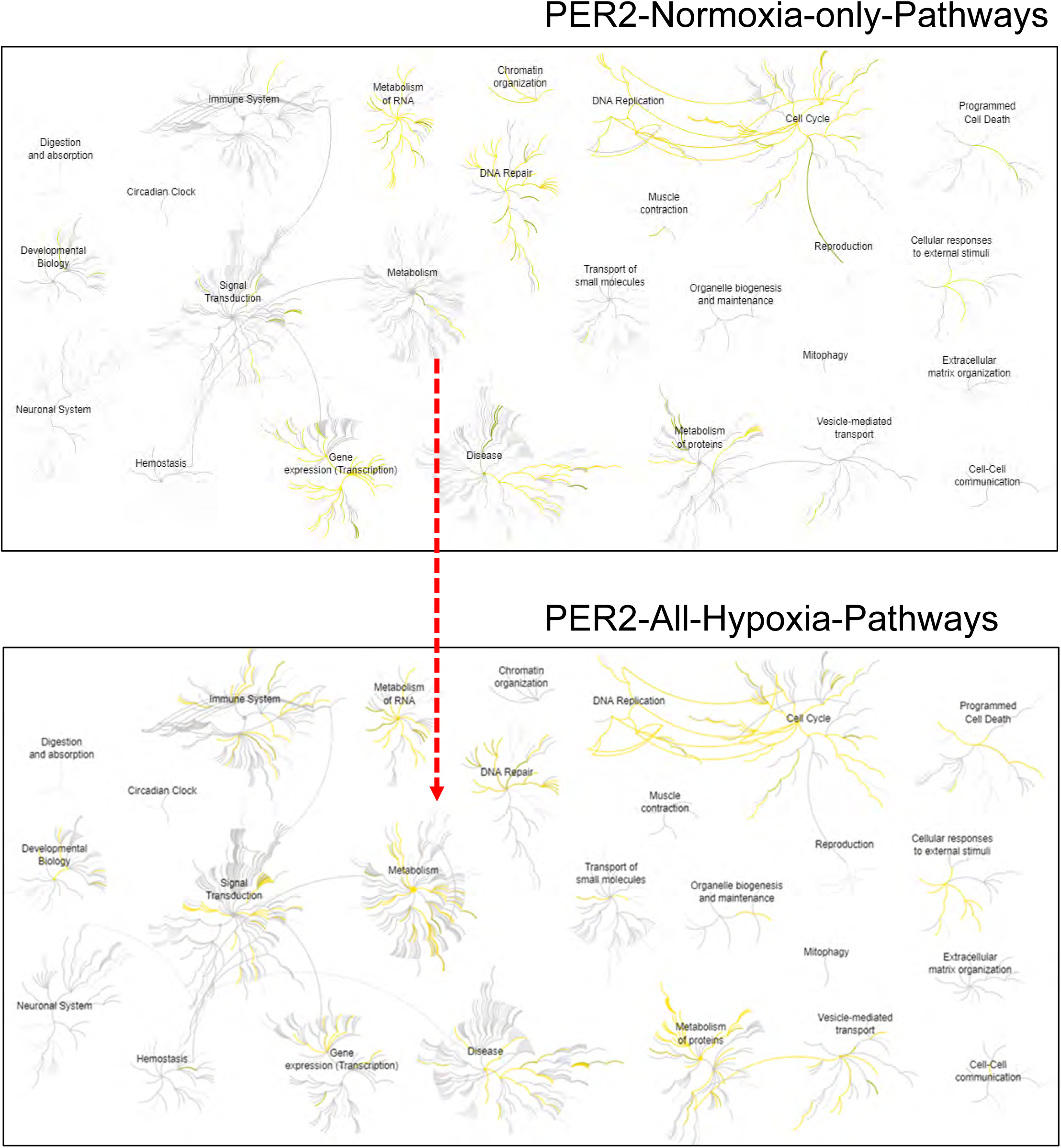
PER2-Normoxia-Hypoxia-Pathways*. Reactome* analysis of an affinity purification–mass spectrometry-based proteomics from hypoxic HMEC-1 cells, indicating as strong involvement of PER2 in metabolic pathways under hypoxia. Yellow depicts PER2 pathways in comparison to all available *Reactome* pathways (grey).

**Figure S6. Related to Figure 4.**
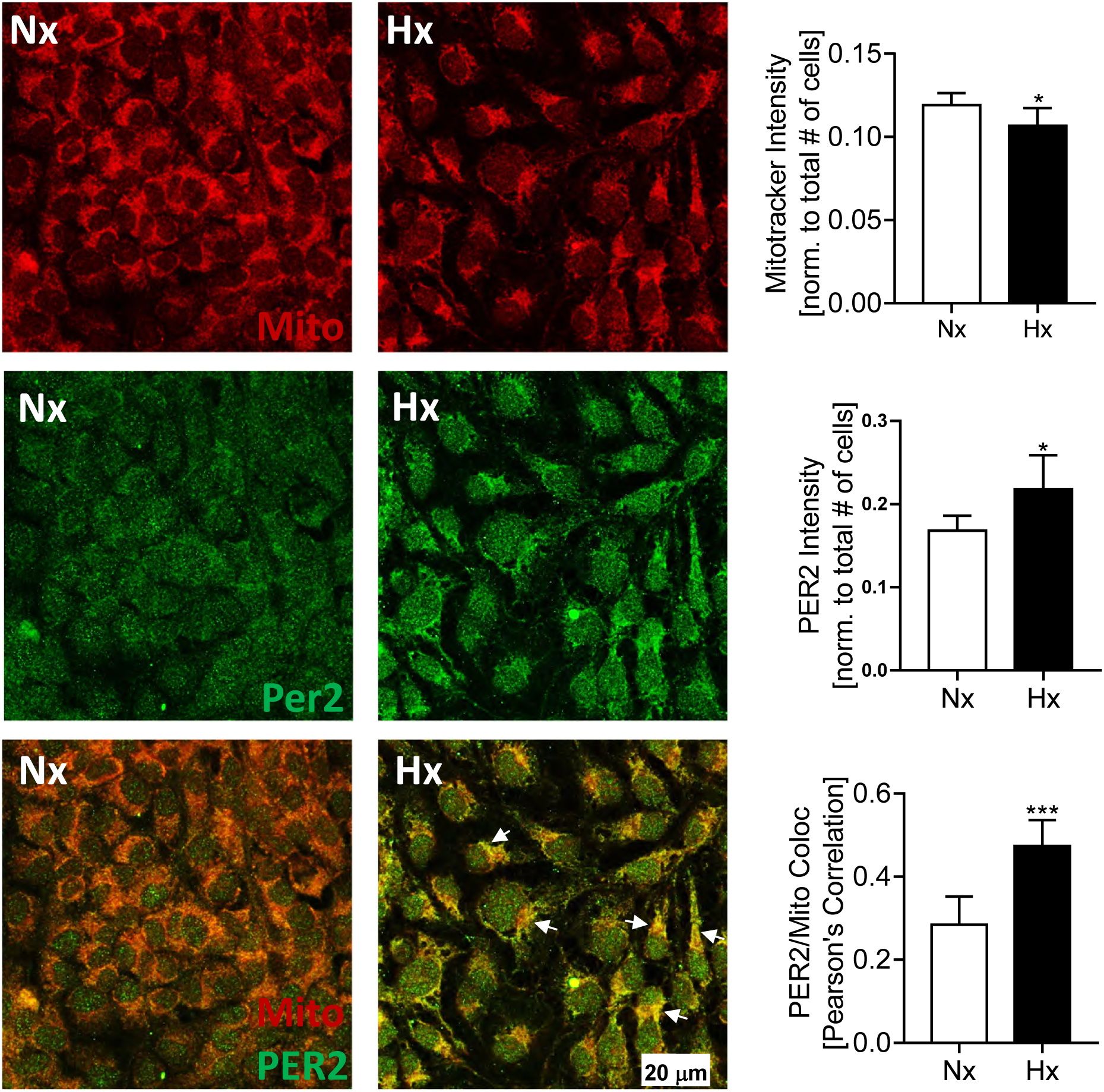
Colocalization of PER2 and mitochondria (upper panel: MitoTracker Red CMXRos staining (red), middle panel: PER2 staining (green), lower panel: overlay MitoTracker Red and PER2 staining). Shown are staining from HMEC-1 cells after 24h of normoxia or 24h of hypoxia 1%. White arrows indicate mitochdrial translocation of PER2 (yellow, mean±SD, n=6, *=*P*<0.05, ***= *P*<0.001).

**Figure S7. Related to Figure 4 and Figure 5.**
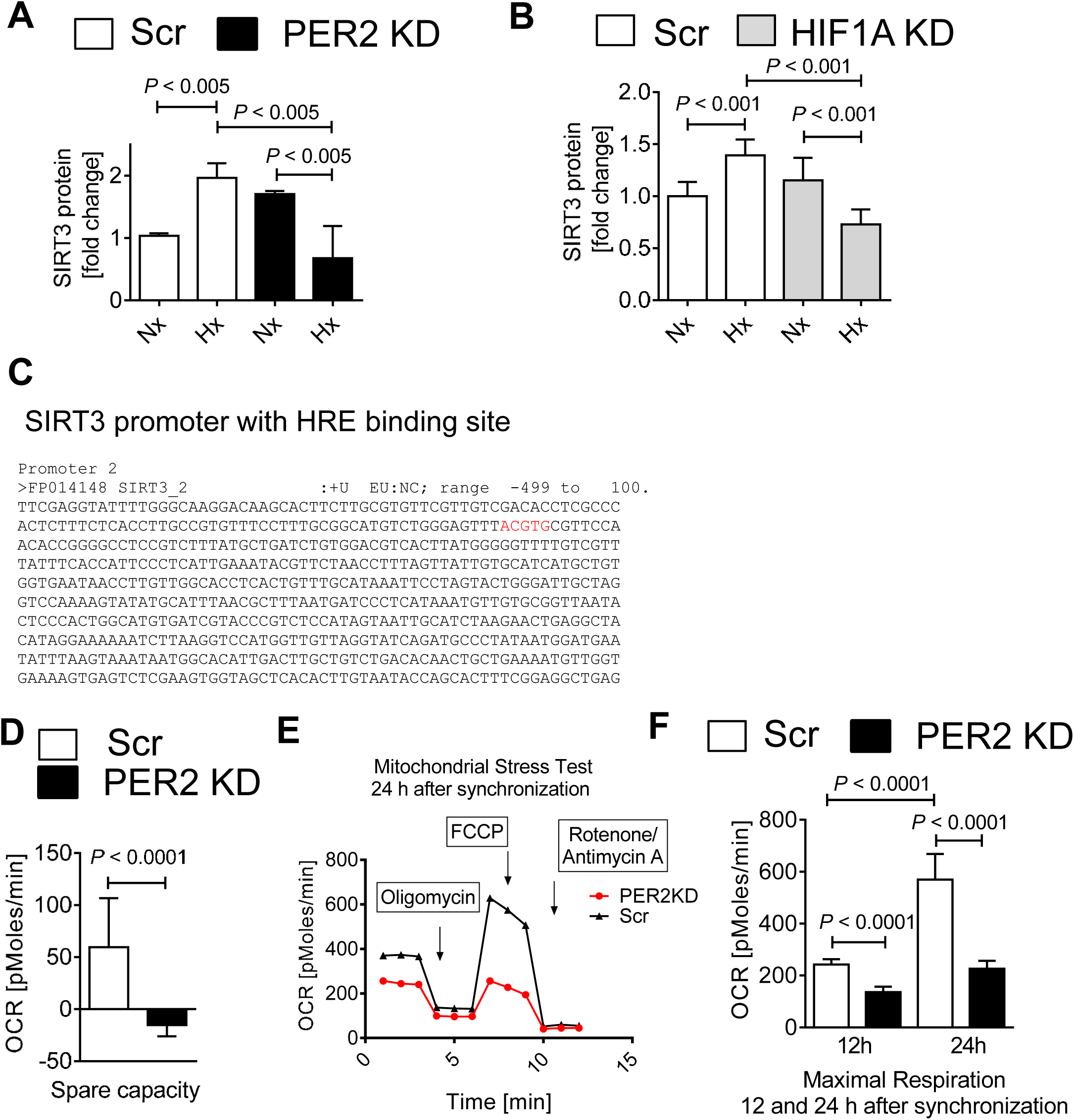
(**A**) Quantification of SIRT3 protein from PER2 KD or Scr control HMEC-1 in Hx (hypoxia 1%) or Nx (normoxia; mean±SD, n=5). (**B**) Quantification of SIRT3 protein from HIF1A KD or Scr control HMEC-1s in Hx or Nx (mean±SD, n=5). (**C**) Region of the human SIRT3 promoter containing a hypoxia response element (HRE) binding site (red). (**D**) Mitochondrial stress test in PER2 KD or Scr control HMEC-1 measuring spare capacity 12 h after cell synchronization (mean±SD, n=10). (**E, F**) Mitochondrial stress test administered at 24h after cell synchronization in PER2 KD or Scr control cells (mean±SD, n=5). Differences in maximal respiration between time point 12 and 24h are quantified in (**F**).

**Figure S8. Related to Figure 5.**
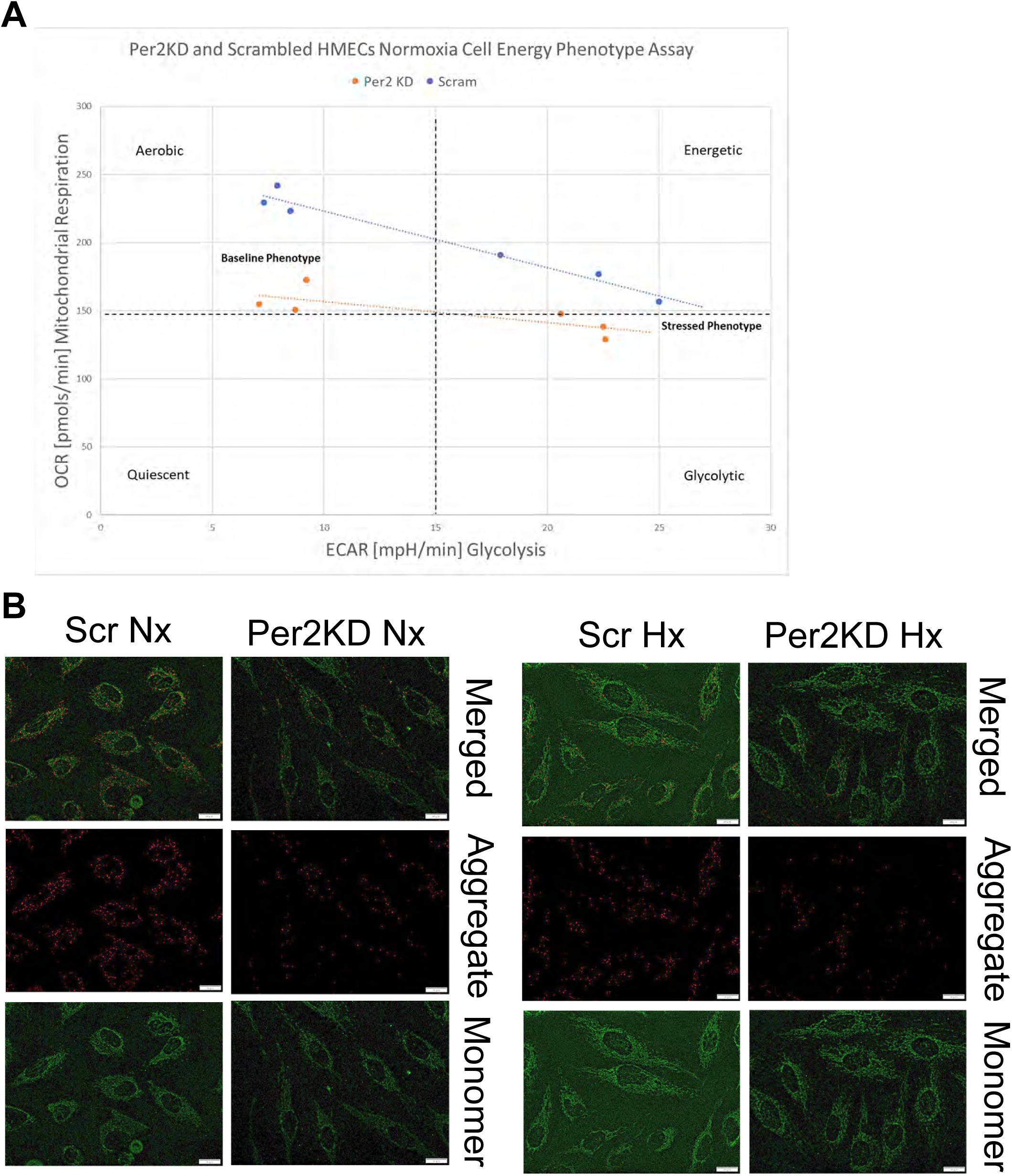
(**A**) Cell energy phenotype test using the Seahorse Bioanalyzer in PER2 KD or Scram control HMEC-1 at baseline. Quiescent phenotype = cell not energetic for either metabolic pathway; energetic phenotype = cell uses both metabolic pathways; aerobic phenotype = cell uses predominantly mitochondrial respiration; and glycolytic = cell uses predominantly glycolysis (mean±SD, n=10). (**B**) JC-1 staining results from PER2 KD or Scr control HMEC-1s in Hx (hypoxia, 1%) or Nx (normoxia). Aggregate represents hyperpolarized cells and monomer represents depolarized cells (mean±SD, n=3, white scale bar=20μm).

**Figure S9. Related to Figure 5.**
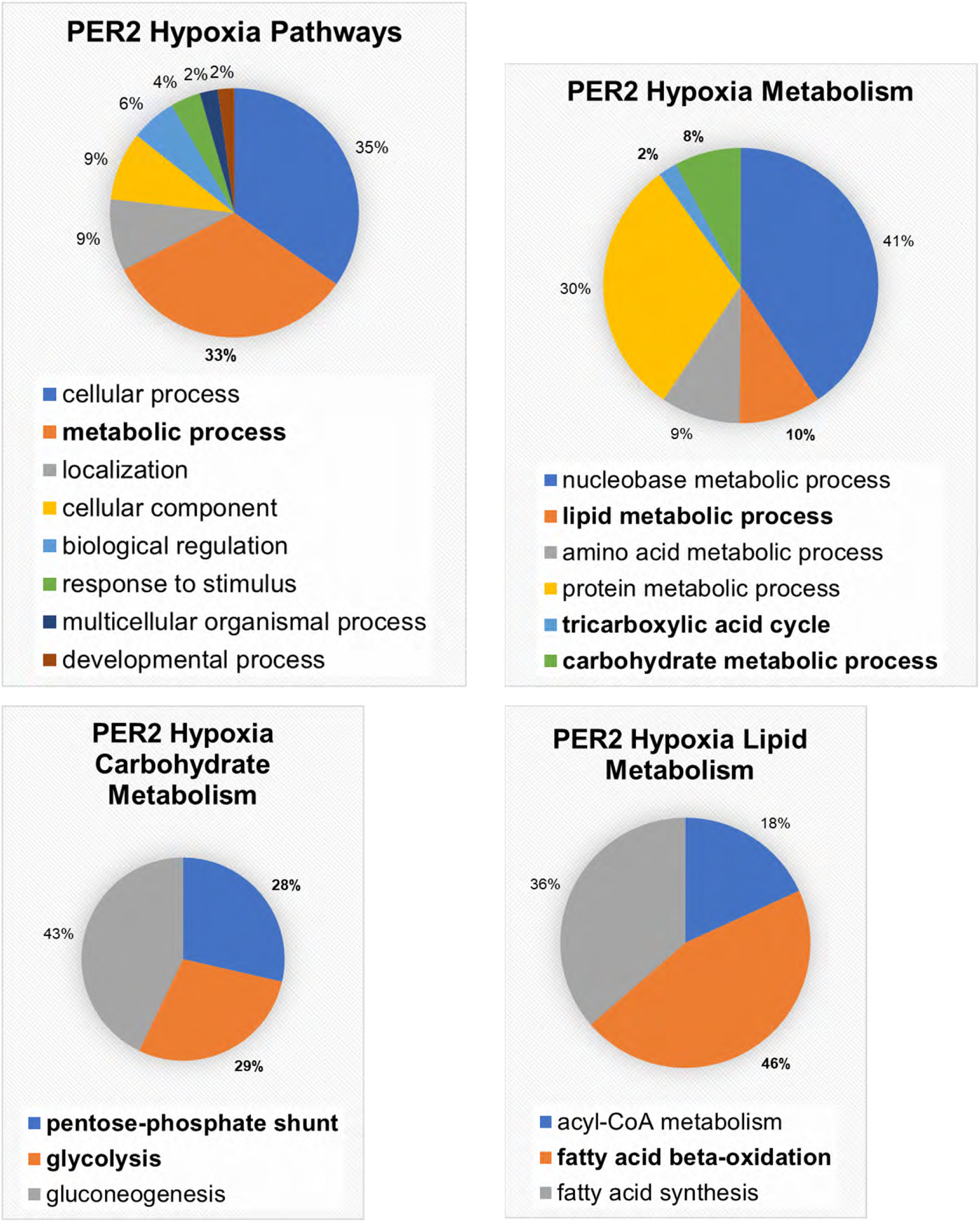
PER2-Hypoxia-only-Pathways. PANTHER (Protein ANalysis THrough Evolutionary Relationships) analysis of an affinity purification–mass spectrometry-based proteomics from hypoxic HMEC-1 cells.

**Figure S10. Related to Figure 7.**
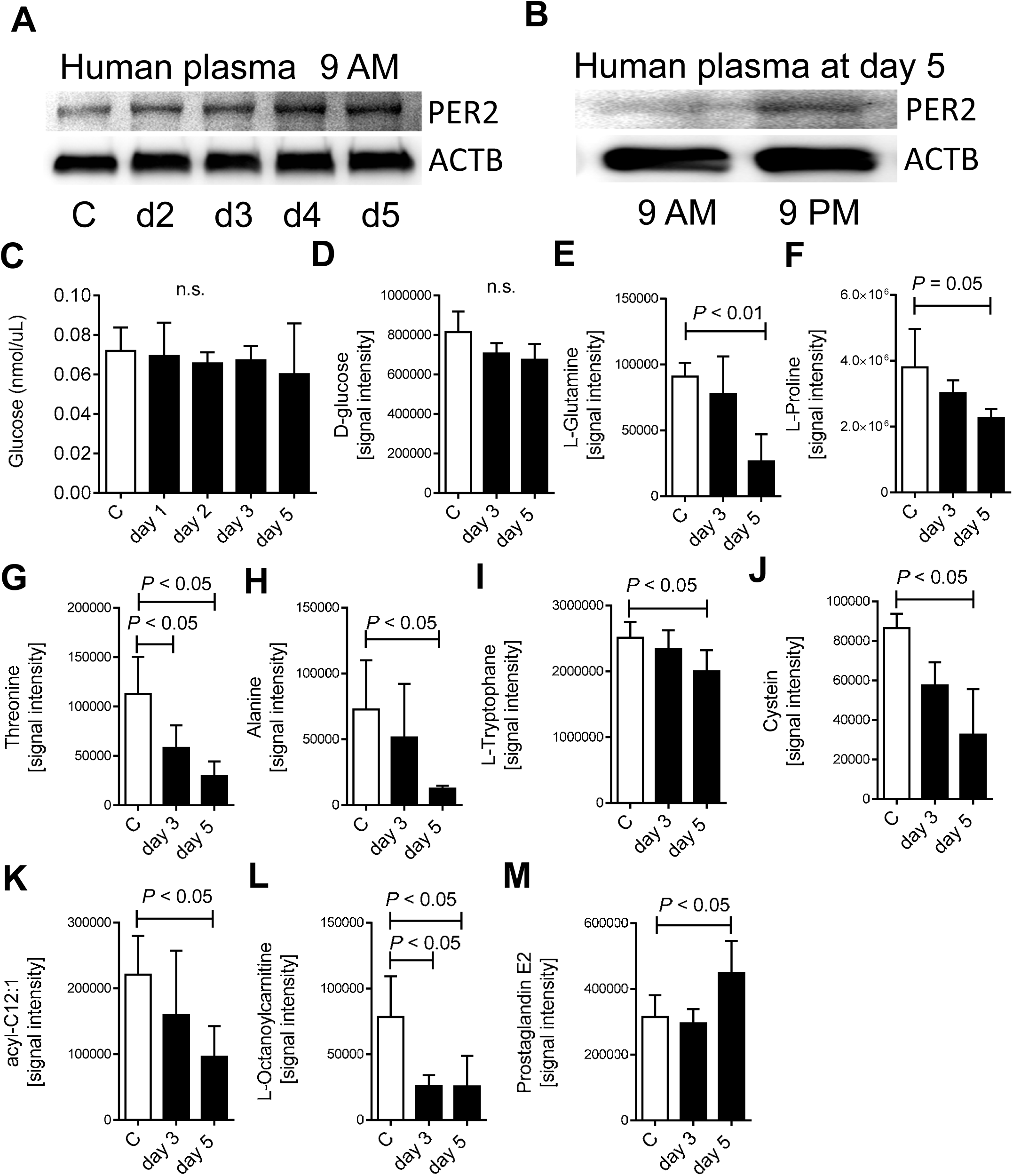
(**A, B**) Immunoblots of plasma PER2 at 9AM and 9PM from one healthy human volunteer exposed to 5 days of intense light for 30 min between 8.30 and 9.00 AM. (**C**) Plasma glucose levels from human healthy volunteers during 5 days of intense light therapy. (**D-M**)Targeted metabolomics in plasma samples from healthy human volunteers exposed to 5 days of 30 minutes intense light from 8.30 to 9.00 AM each morning; (mean±SD; n=3).

**Figure S11. Related to Figure 7.**
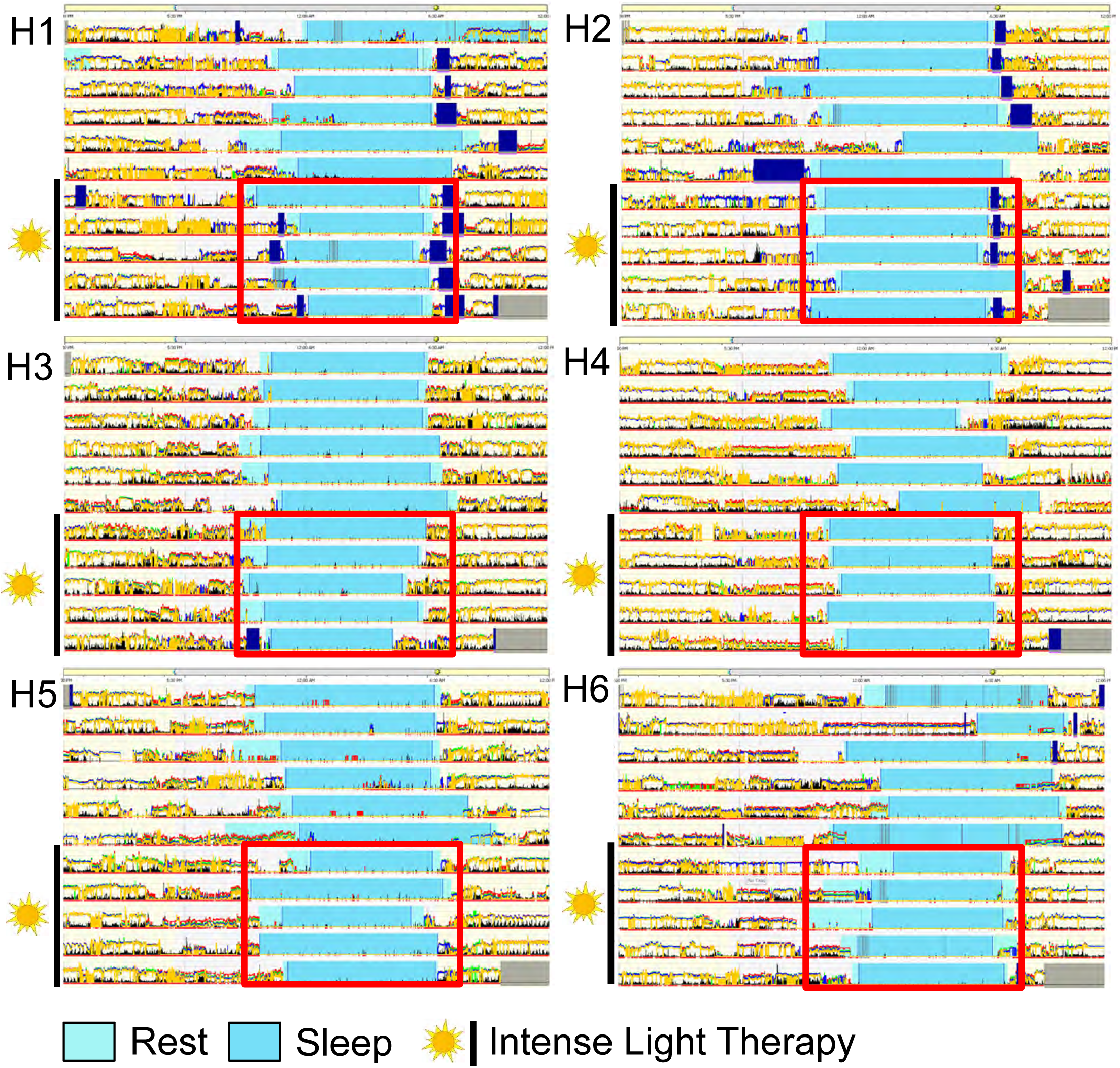
Actigraphy data using a validated accelerometer (Actiwatch 2) from human healthy volunteers during 5 days without and 5 days with intense light therapy (30 min intense light from 8.30 – 9.00 AM; n=6, H=healthy volunteer; **Note**: synchronized sleep phases during intense light exposure [red square] vs no intense light therapy.

